# Functional brain region-specific neural spheroids for modeling neurological diseases and therapeutics screening

**DOI:** 10.1101/2022.05.04.490442

**Authors:** Caroline E. Strong, Srikanya Kundu, Molly Boutin, Yu-Chi Chen, Kelli Wilson, Emily Lee, Marc Ferrer

**Author notes:** Corresponding Authors: Marc Ferrer, Division of Preclinical Innovation, National Center for Advancing Translational Sciences, National Institutes of Health, 9800 Medical Center Dr, Building B, Rockville, MD, 20850, Emily Lee, Division of Preclinical Innovation, National Center for Advancing Translational Sciences, National Institutes of Health, 9800 Medical Center Dr, Building B, Rockville, MD, 20850.

## Abstract

3D spheroids have emerged as powerful drug discovery tools given their high-throughput screening (HTS) compatibility. Here, we established a method for generating functional neural spheroids with differentiated human induced pluripotent stem cell (hiPSC)-derived neurons and astrocytes at cell type compositions mimicking specific regions of the human brain. Recordings of intracellular calcium oscillations were used as functional assays, and the utility of this spheroids system was shown through disease modelling, drug testing, and formation of assembloids to model neurocircuitry. We developed disease models for Alzheimer’s and Parkinson’s Disease through incorporating genetically engineered cells into spheroids, and Opioid Use Disorder by chronically treating spheroids with DAMGO. We reversed baseline functional deficits in each disease model with clinically approved treatments and showed that assembloid activity can be chemogenetically manipulated. Here, we lay the groundwork for brain region-specific spheroids as a robust functional assay platform for HTS studies.

## INTRODUCTION

Therapeutic development for neurological diseases has been hindered by a lack of predictable *in vitro* cellular assays, limited *in vivo* animal models, and high cost and complexity of clinical trials (DiMasi et al., 2016; Wong et al., 2019). As a result, less than 10% of clinical candidates to treat neurological diseases are approved by regulatory agencies each year (DiMasi et al., 2016; Hay et al., 2014; Wong et al., 2019). The increase in neurodegenerative disease occurrence and substance use disorders over the last few decades makes it critical to develop technologies to improve the success rate of drug discovery for neurological diseases (Hou et al., 2019; Blanco et al., 2021; Rehm and Shield, 2019). One way to improve the predictability of assay platforms for drug discovery is to develop functional neural cellular models with high physiological and pathological relevance.

A range of iPSC-derived neural models have been developed, including 2D monolayer cultures and 3D spheroids and organoids, which include different neuronal types that may include some brain-like morphology (Di Lullo and Kriegstein, 2017; Pasca, 2018). While 2D neural models are compatible with high-throughput screening (HTS) assays, neural organoids are better at recapitulating *in vivo* morphology (Pasca, 2018). While organoids acquire greater cellular diversity and some brain-like organization, their complexity, batch-to-batch variation, limited co-culture differentiation, and lengthy maturation times can limit their use for HTS (Andrews and Kriegstein, 2022). Cortical neuronal spheroids, which have been produced from neural stem cells (NSCs) that mature into differentiated excitatory and inhibitory neurons in the spheres (Dingle et al., 2015; Mariani et al., 2015), may be a more readily adaptable model for HTS than organoids, but are currently mostly limited to cortical neurons, thus limiting the cell type composition. The lack of more diverse neuronal subtype populations such as dopaminergic neurons restricts the applications of these spheroids for disease modeling (Mariani et al., 2015; Nzou et al., 2018; Pasca et al., 2015; Woodruff et al., 2020; Yoon et al., 2019).

Here, we describe the development of an HTS-compatible, functional neural spheroid system, referred to here as brain-region specific spheroids, that can mimic the physiology of distinct brain regions, including the prefrontal cortex (PFC) and ventral tegmental area (VTA). In addition to healthy neural spheroid models, we also describe the development of Alzheimer’s Disease (AD) and Parkinson’s Disease (PD) neuronal spheroid models using iPSC-derived neurons with genetically engineered, disease-relevant mutations, as well as an Opioid Use Disorder (OUD) neuronal model induced by chronic treatment with a µ-opioid receptor agonist. We measured intracellular calcium oscillations as an HT-compatible, functional readout of neural activity, as calcium oscillations haves been previously shown to be highly correlated with the electrophysiological properties of neurons (Ali et al., 2020), and observed phenotypic differences when compared to “wildtype”, or healthy, controls with the mutant, diseased spheroids. A machine learning classifier model was used to quantify phenotype labeling predictability and showed high accuracy for both the AD and PD models (>94%). Furthermore, clinically approved treatments for each disease were used to treat spheroids, and a reversal of deficits was observed for all three disease phenotypes. Finally, to address whether we could also extend this technology to neural circuit-specific modeling, we created functional assembloids using fused VTA- and PFC-like spheroids, and using chemogenetic modulation, we were able to demonstrate assembloid plasticity. Together, this study establishes the utility of these brain region-specific neural spheroids through disease and neurocircuitry modeling, and for use as HTS-compatible drug screening platforms.

## MATERIALS AND METHODS

### Generation and maintenance of designer spheroids

#### Cells and Donor Information

Matured, differentiated iPSC-derived cells were obtained from FujiFilm CDI. Wildtype (Wt) cells included iCell DopaNeurons (cat #R1088), iCell GlutaNeurons (cat #R1061), iCell GABANeurons (cat #R1013) and iCell Astrocytes (cat #R1092). iCell Dopa Neurons PD SNCA A53T HZ (cat # R1109) were used to model Parkinson’s Disease (PD) while iCell GABANeurons (APOE e4/4) (cat# R1168) were used to model Alzheimer’s Disease. The donor ID for Wt and A53T iCell DopaNeurons and iCell GlutaNeurons was 01279, a healthy male age 50-59. A53T iCell DopaNeurons were genetically engineered to have the SNCA A53T mutation. The donor ID for iCell Astrocytes as well as Wt and APOE iCell GABANeurons was 01434, a healthy female <18 years old. The APOE e4/4 line was genetically modified to encode the APOE e4/4 mutation.

#### Tissue Culture Media

After each cell type was thawed, base media with supplements was used to create a cell suspension. Base media used to form spheroids differed by cell type; iCell Base Medium 1 (CDI, #M1010) was used for iCell DopaNeurons, iCell GABANeurons, and iCell Astrocytes while BrainPhys Neuronal Medium (Stem Cell Technologies, #05790) was used for iCell GlutaNeurons. Supplements and iCell Base Medium 1 were provided in the iCell kits referenced above, and 2% Neural Supplement B plus 1% Nervous System Supplement were added to media for iCell DopaNeurons, while media for iCell GABANeurons contained 2% Neural Supplement A. Media for iCell GlutaNeurons contained 2% Neural Supplement B, 1% Nervous System Supplement, 1% N2 supplement (Thermo, 17502048), and 0.1% laminin (Invitrogen, #23017-015) in BrainPhys media. Base media for iCell GABANeurons was used for Astrocytes. 45ul of cell suspension was used to seed each well of a 384 well, ULA, round bottom plate (Corning, Cat # 3830), and plates incubated overnight at 37C, 5% CO2 incubator.

The day after spheroids were plated, maintenance media was added such that each well contained 90 uL with 45 uL consisting of base media and 45 uL consisting of maintenance media. Maintenance media was the same for all spheroids and consisted of BrainPhys Neuronal Medium (Stem Cell Technologies, cat #05790) supplemented with 1X N2, 1X B27 (Thermo, cat# 17502048 & 17504), 20 ng/mL BDNF and 20 ng/mL GDNF (Stem Cell Technologies, cat# 78005 and 78058, respectively), 1 ug/mL laminin (Invitrogen), 1 mM cAMP (Tocris, cat# 1141), and 20 nM ascorbic acid (Tocris, cat# 4055). A stock solution consisting of all materials except cAMP and ascorbic acid was prepared in advance, and the cAMP and ascorbic acid were added to the media fresh on each day of media changes. Half media changes occurred every other day, and spheroids were maintained for 3-weeks at 37C, 5% CO2.

#### Cell Thawing

For thawing, iCell DopaNeurons (“Dopa”), iCell GABANeurons (“GABA”), and iCell Astrocytes (“Astro”) were placed in a 37C water bath for 3 min and iCell GlutaNeurons (“Gluta”) neurons for 2 min, according to the manufacturer’s instructions. The contents of each vial were dispensed into separate 15 mL conical tubes. Base media (1 mL) for each cell type was added to the empty cell vials to collect any remaining cells, dispensed in drop-wise fashion on top of the cell suspension in each tube, then 8 mL of media was added to each tube. Tubes containing cell suspension of GABA and Astro cells were centrifuged at 300 g x 5 min, while tubes with either Dopa or Gluta cell suspension were centrifuged at 400 g x 5 min. The supernatant was aspirated and resuspended in 2 mL of base media, then cells for each cell type were counted using a Countess Cell Counter (Thermo).

#### Generation of spheroids

After counting, base media was added to achieve a cell suspension containing 5e5 cells/mL for each cell type. Cell types required in each spheroid type were then mixed in fresh 50 mL conical tubes. The cell type compositions of control spheroids include: 100% dopa, 100% gluta, 100% GABA, 90% dopa + 10% astro, 90% gluta + 10% astro, 90% GABA + 10% astro. A proof-of-concept study (Fig. 2) generated 16 types of spheroids consisting of 90% neuron and 10% astrocyte but with differing percentages of neuronal subtypes, and these spheroid compositions can be found in the figures. Brain region-specific spheroids modeling the ventral tegmental area (VTA-like) and prefrontal cortex (PFC-like) each contained 90% neurons + 10% astrocytes with differing neuronal cell type compositions. VTA-like spheroids contained 65% dopa, 5% gluta, and 30% GABA neurons while PFC-like spheroids contained 70% gluta + 30% GABA neurons. Once a cell suspension was created containing the cell type composition needed, 50 uL was manually dispensed with a 16-channel multichannel Finnpipette (Thermo) into 384-well round bottom ultra-low attachment (ULA) spheroid microplates (Corning, cat# 3830). Plates were then sealed with parafilm and centrifuged at 1485 x g for 10 min to pull cells to the bottom of the plate. One day later, 5 uL of base media was removed and 45 uL maintenance media was added as described above.

#### Generation of assembloids

Assembloids were formed with VTA- and PFC-like spheroids such that one spheroid type was transduced with AAV9-GCaMP6f (Addgene, cat# 100836-AAV9) while the other was transduced with either an inhibitory or excitatory DREADDs virus (Addgene, cat# 50475-AAVrg and 50474-AAVrg). Assembloids were made 1-week prior to recording by placing one of each spheroid type into a 1.7 mL tube together. Both spheroids were pulled up into a wide bore 200 uL pipette tip (Rainin, cat# 30389188) with only 15 uL media and dispensed into the bottom of a well int the Corning ULA round bottom plates (#3830). Collagen I (Fisher, cat# CB354249) was made with media, 10X phosphate buffered saline, and 1 N NaOH at a 3 mg/mL concentration and 15 uL was pipetted on top of the two spheroids. The plate was centrifuged at 1485 g x 2-min and placed in the incubator at 37C for overnight gelling and the following day, 50 uL media was added to the wells. Maintenance media used was the same as spheroids and half media changes were performed every other day prior to testing.

### Measurement of calcium activity

#### Viruses and dyes

To assess calcium activity, the calcium dye, Cal6 (Molecular Devices), and genetically encoded calcium indicator, GCaMP6f (Addgene, cat# 100836-AAV9) were used. Cal6 was used according to manufacturer’s instructions and 10 mL of maintenance media was added to each vial. Two hours before activity was recorded, half of the spheroid media was exchanged for media with Cal6. The plates were covered in foil and placed in the incubator at 37C during the 2-hr incubation period. Adeno-associated viruses (AAV) were added to the media on day 7 to allow for 2-weeks expression prior to recording or testing at 2e5 multiplicity of infection (MOI). GCaMP6f infection occurred via an adeno-associated virus serotype 9 (AAV9) expressed under the CAG promoter for expression in both neurons and astrocytes. Designer receptors exclusively activated by designer drugs (DREADDs) viruses were used to stimulate and inhibit neuronal activity within spheroids. Both viruses were retrograde AAVs expressed under the human Synapsin promoter for expression in neurons and fused with an mCherry fluorophore. The DREADDs viruses either inserted the designer receptor hM4D(Gi), an inhibitory G-protein coupled receptor, or hM3D(Gq), a stimulatory G-protein coupled receptor (custom made Chemogenetics AAV: Addgene). Clozapine-N-oxide (CNO, Tocris cat# 4936) was suspended in dimethyl sulfoxide (DMSO) and used as the designer drug to activate the DREADDs viruses. CNO was tested at 1 and 10 μM, with data reported from the 1 μM concentration.

#### Fluorescent Imaging plate Reader (FLIPR) measurements

The FLIPR Penta (Molecular Devices) was used to assess calcium fluorescence across the 384-well plate simultaneously and to observe changes after treatment with compounds. The evening before calcium activity was measured, plates were centrifuged at 1485 x g for 2-min to get spheroids to the bottom of the well in a centered position. On the day of recording, the plate used for recording was placed in the read plate position inside of the FLIPR following the 2-hr Cal6 incubation at 37C. Standard filter sets were used for Cal6 imaging with excitation set at 470-495 and emission at 515-575 nm. Fluorescent image reads were taken every 0.6 seconds for all plates, with exposure time of 0.03 and 50% excitation intensity. Recordings from the initial seven plates consisted of 1000 reads and were 10-min recordings with 2.5 gain, while the final two plates consisted of 500 reads (5-min recordings) with a gain of 2. Baseline recordings were taken across all plates and, if applicable, more recordings were obtained 1-, 30-, 60-, and/or 90-min after compound treatment. In between recordings, plates were wrapped in foil and placed back in the incubator at 37C, 5% CO2.

#### Confocal Imaging

The Opera Phenix Plus High-Content Imaging System (Perkin Elmer) spinning disk confocal was used to record calcium activity from spheroids in individual wells. Prior to recording, the stage was pre-warmed to 37C, and the carbon dioxide was set to 5%. Recordings were obtained both from spheroids expressing GCaMP6f along with those incubated in Cal6 dye. Recordings were captured with a 20X water immersion objective 55um from the bottom of the well, and were obtained at a frame rate of 1.6 frames/sec with 480 frames total, making the recordings 5-min. The protocol was set to record well to well such that recordings were automated but taken from one spheroid at a time before recording from subsequent wells. For all calcium activity recordings obtained from the Phenix Plus, the FITC channel was used where excitation was set to 488 nm and emission at 535 nm. For spheroids expressing GCaMP6f, the exposure time was set to 20 ms and the laser power was set to 30% while for spheroids in Cal6 dye, the exposure time was 20 ms with laser power set to 10%. The focal plane for recordings from assembloids varied depending on where they were suspended in the collagen, though these recordings were all obtained within 250 µm from the bottom of the well. For assembloids, the exposure time was set to 40 ms with laser power set to 40%.

### Drug Treatment

#### Chronic DAMGO treatment

To model opioid use disorder (OUD), a subset of spheroids was treated with DAMGO (Tocris, cat# 1171), a selective mu opioid receptor (MOR) agonist, chronically during the 3-week spheroid maintenance period. DAMGO was reconstituted in water at a 1 mM concentration and diluted in media to 20 uM for a final concentration of 10 uM after the half media exchange. For spheroids subjected to chronic DAMGO treatment, 20 uM DAMGO in maintenance media was added via half media exchange beginning on day 10, with treatments occurring every other day for 10 days, giving a total of five treatments. For spheroids subjected to DAMGO withdrawal, the same protocol was followed except that spheroid did not receive DAMGO for the final treatment and instead were subjected to a three day washout period intended to model the withdrawal aspect of OUD.

#### 384 Pin Tool compound addition

A 384 well pin tool (Rexroth) was used to transfer compounds simultaneously to the spheroid plate. Compound transfer via the 384 well pin tool occurred after the baseline recordings with the fluorescent imaging plate reader (FLIPR). 60 nL of compound suspended in DMSO was transferred to 60 uL of media in each well. Immediately after compound transfer, the spheroid plate was either placed back inside of the FLIPR for a recording 1-min after compound treatment or placed back in the incubator if post-treatment recording was >30-min after compound transfer.

### Cell viability assays

#### 3D Cell Titer Glo

CellTiter-Glo 3D Cell Viability Assay (Promega, cat# G9681) was used according to the manufacturer’s instructions. CellTiter-Glo 3D Reagent was thawed overnight at 4C and brought to room temperature (RT) for 20-min before use. After the FLIPR assay, 30 uL of CellTiter-Glo 3D Reagent was added to the spheroid plate and was mixed by shaking for 5-min at RT followed by a 25-min incubation period off the shaker at RT. Luminescence was read using a PHERAstar FSX microplate reader (BMG LabTech) to measure amount of ATP present.

#### Calcein and Propidium Iodide (PI) staining

Imaging of live and dead cells was done via Calcein (Thermo, cat# C1430) and PI (Thermo, cat# P3566) staining on live spheroids. Calcein AM and PI were diluted in 1X Dulbecco’s phosphate-buffered saline (DPBS; Thermo, cat# 14040141) to concentrations of 1:2000 and 1:1000 to achieve final concentrations of 0.5 and 1 uM, respectively. Half of the media was removed from each well (45 uL) and was exchanged with Calcein AM and PI in DPBS. Spheroids were incubated at 37C for 30-min prior to live cell imaging. For imaging, spheroids were placed in the Phenix Plus with the stage pre-warmed to 37C and 5% carbon dioxide circulating. The FITC channel, with 488 nm excitation and 535 nm emission, was used to image Calcein AM while Cy3 (excitation of 530 nm, emission of 620 nm) was used to image PI. 150 uM image stacks were collected with a 10X air objective using a 2 um z-step.

### Tissue Processing

#### Spheroid fixation

Spheroids were fixed with 4% paraformaldehyde (PFA) in PBS overnight at 4C. The following day, spheroids were washed with PBS, where half of the PFA was removed and exchanged with PBS, a total of four times. On the final wash, PBS with 0.1% sodium azide (Sigma, cat# S2002) was added for spheroid preservation. Plates were sealed with parafilm and stored at 4C until further use.

#### Immunohistochemistry (IHC)

IHC was used to stain for neurons and astrocytes, neuronal cell type markers, and pre- and postsynaptic markers. For neurons, polyclonal chicken anti-MAP2 was used while astrocytes were stained with rabbit polyclonal anti-GFAP antibody (Abcam, cat# ab5392, ab7260). For neuronal cell type markers, polyclonal chicken anti-Tyrosine Hydroxylase (TH) was used to stain dopaminergic neurons, monoclonal mouse anti-vGluT1 was used to stain glutamatergic neurons, and polyclonal rabbit anti-Parvalbumin (PV) was used to stain GABAergic neurons (Abcam, cat# ab76442, ab242204, ab11427). Mouse monoclonal anti-bassoon antibody was used as a presynaptic marker while rabbit polyclonal anti-homer1 antibody was used as a postsynaptic marker (Abcam, cat# ab82958, ab97593). For the immunostaining assay, all liquid removal steps were performed via manual pipetting and all incubation steps occurred on a shaker. PBS with 0.1% azide was removed and blocking solution consisting of 5% normal goat serum (NGS), 2% bovine serum albumin (BSA; Fisher, cat# BP1605), and 0.5% Triton X-100 (Sigma, cat# X100) in PBS was added for 30-min. After 30-min, half of the blocking solution was removed and primary antibodies made in blocking solution were added at double the desired concentration for final concentrations and times as follows: Anti-MAP2 and anti-GFAP 1:500, overnight at 37C on shaker; anti-Homer 1:100, anti-bassoon 1:100. 72 hours, 37C, on shaker. Spheroids were then washed with PBS + 0.3% triton X-100 (PBT) in three half changes followed by an additional three washes with 15-min incubations. Secondary antibodies at final staining concentration of 1:600 in blocking buffer were incubated at 37C overnight, shaking, at concentrations as follows: goat anti-chicken Alexa Fluor 647, goat anti rabbit Alexa Fluor 488, and goat anti-mouse Alexa Fluor 647 (Invitrogen, cat# A32933, A32731, A32728, respectively).Hoescht was used at 1:10,000 for nuclei staining and washes occurred the same as described above with primary antibodies.

#### Fluorescent *in situ* Hybridization (FISH)

FISH was performed to validate glutamatergic cell type compositions in brain region specific designer spheroids modeling the VTA and PFC. Homo sapiens solute carrier family 17 (vesicular glutamate transporter) member 7 mRNA (Hs-SLC17A7, cat# 415611; GenBank Accession Number: NM_020309.3) was used to probe for glutamatergic neurons according to the RNAScope Multiplex Fluorescent Reagent Kit v2 user manual (Advanced Cell Diagnostics, cat# 323100). Spheroids fixed in 4% PFA were used and therefore the tissue was pre-treated according to the formalin-fixed paraffin-embedded (FFPE) preparation method, though the deparaffinizing step was skipped since spheroids were not paraffin-embedded. After spheroids were incubated in hydrogen peroxide, they were washed with DI water and incubated in protease plus for 30-min at 40C. Protease plus was washed out with DI water and the vGluT was assigned to channel 2 (SLC17A7-C2). After hybridization of the probes, pre-amplification and amplification reagents were applied according to the user guide, where AMP1 was added for 30-min at 40C followed by AMP2 was added for 30-min at 40C and AMP3 was added for 15-min at 40C. The fluorescent Opal 570 dye was added to channel 2 containing SLC17A7-C2 (excitation: 550 nm, emission: 570 nm; Akoya Biosciences). DAPI was added in the final step to spheroids for 30-sec, then washed with PBS. Spheroids remained in PBS until tissue clearing reagent was added. Tissue was washed in 1X wash buffer twice for 2-min each time between incubations after probe hybridization steps. Prior to probe hybridization, spheroids were washed with DI water in accordance with the manufacturer’s instructions.

#### Tissue Clearing

After immunostaining or FISH, ScaleS4 Tissue clearing solution was added to spheroids to reduce autofluorescence during image acquisition, as previously described (Hama et al., 2011). ScaleS4 was made with 40% D-sorbitol (Sigma, cat# S6021), 10% glycerol (Sigma, cat# G2289), 4M Urea (Sigma, cat# U5378), 0.2% triton X-100, 15% DMSO (Sigma, cat# D2650) in UltraPure water (Invitrogen, cat# 10977-015). ScaleS4 solution was mixed via shaking at 37C for two days and stored at 4C until future use. Before the clearing solution was added, all PBT for IHC spheroids or all wash buffer for FISH spheroids was removed from the well. For spheroids stained with IHC, the nuclear stain, Hoechst 33342 Thermo, cat# 62249) was added at a 1:2000 dilution in the ScaleS4 clearing solution, and 60 uL of clearing solution plus Hoechst was added to each well. For spheroids that were stained with FISH, no nuclear stain was added since DAPI was added during the assay. Spheroid plates were wrapped in foil and placed on the shaker at 37C overnight. The following day, the plate was sealed with parafilm and placed at 4C until imaging.

### Statistics and Reproducibility

#### Graphical Plots

For time series plots, heatmap plots, correlation matrices and radar plots, Python 3.8 was used. Specifically, the packages matplotlib and seaborn were used to generate radar plots, heatmaps, time series plots, and correlation matrices (Hunter, 2007; Waskom, 2020). The Python packages, pandas and numpy, were used to normalize data prior to graphically plotting (Harris et al., 2020; Reback et al., 2020). For principal component analysis scatter plots, TIBCO Spotfire was used, and for column graphs, GraphPad Prism 9.1 Software was used. Schematics used in Figures 1 and 2 were created with Biorender.

**Figure 1.**
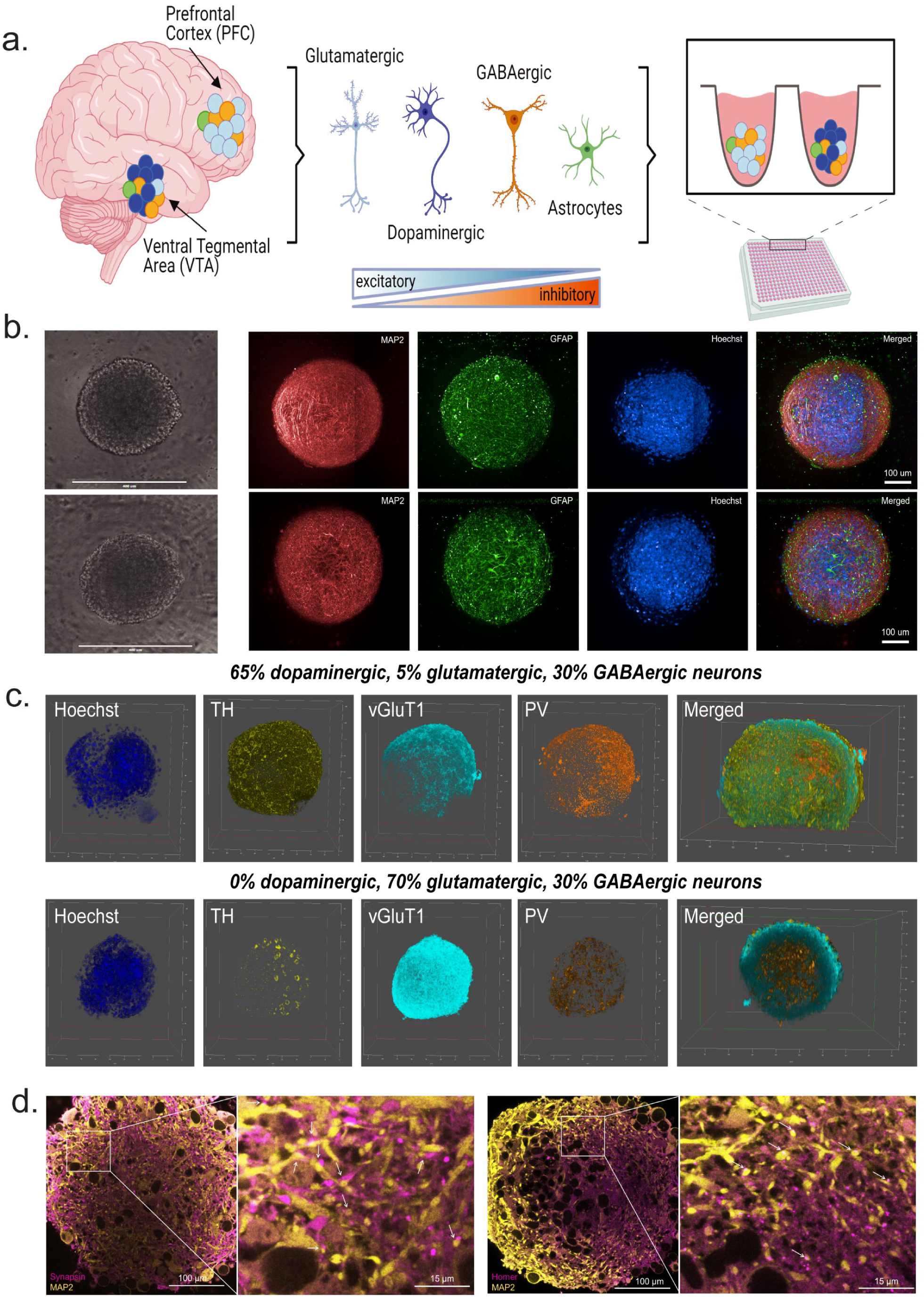
Generation of brain region-specific neural spheroids. **(a.)** Schematic showing method for generating brain region-specific neural spheroids; matured and differentiated iPSC-derived neural cells combined in pre-determined ratios reflecting the cell type compositions of specific regions of the human brain **(b.)** Spheroid types are similar in size regardless of cell type composition (top panel contains mostly dopaminergic neurons, bottom contains mostly glutamatergic neurons; both spheroid types are 90% neuron 10% astrocyte); left to right: brightfield image, MAP2 neuronal marker, GFAP astrocyte marker, Hoechst nuclear stain, merged image of MAP2, GFAP, and Hoechst. 272µm image stack acquired with 0.8 um z-step,20X water objective; represented as maximum projection image of 18 images every 20 slices (16µm). **(c.)** 3D rendering from confocal z-stacks of VTA-like (top panel) and PFC-like (bottom panel) spheroids stained with neuronal cell type markers; Left to right: Hoechst nuclear stain, tyrosine hydroxylase (TH, dopaminergic marker), vGluT1 (glutamatergic marker), and Parvalbumin (PV, GABAergic marker) along with a 3D rendering showing TH, vGluT, and PV expression in the center of each spheroid **(d.)** Confocal images showing MAP2 staining and presynaptic marker, synapsin (left panel), and postsynaptic marker, Homer (right panel), indicating functional synapses.

**Figure 2.**
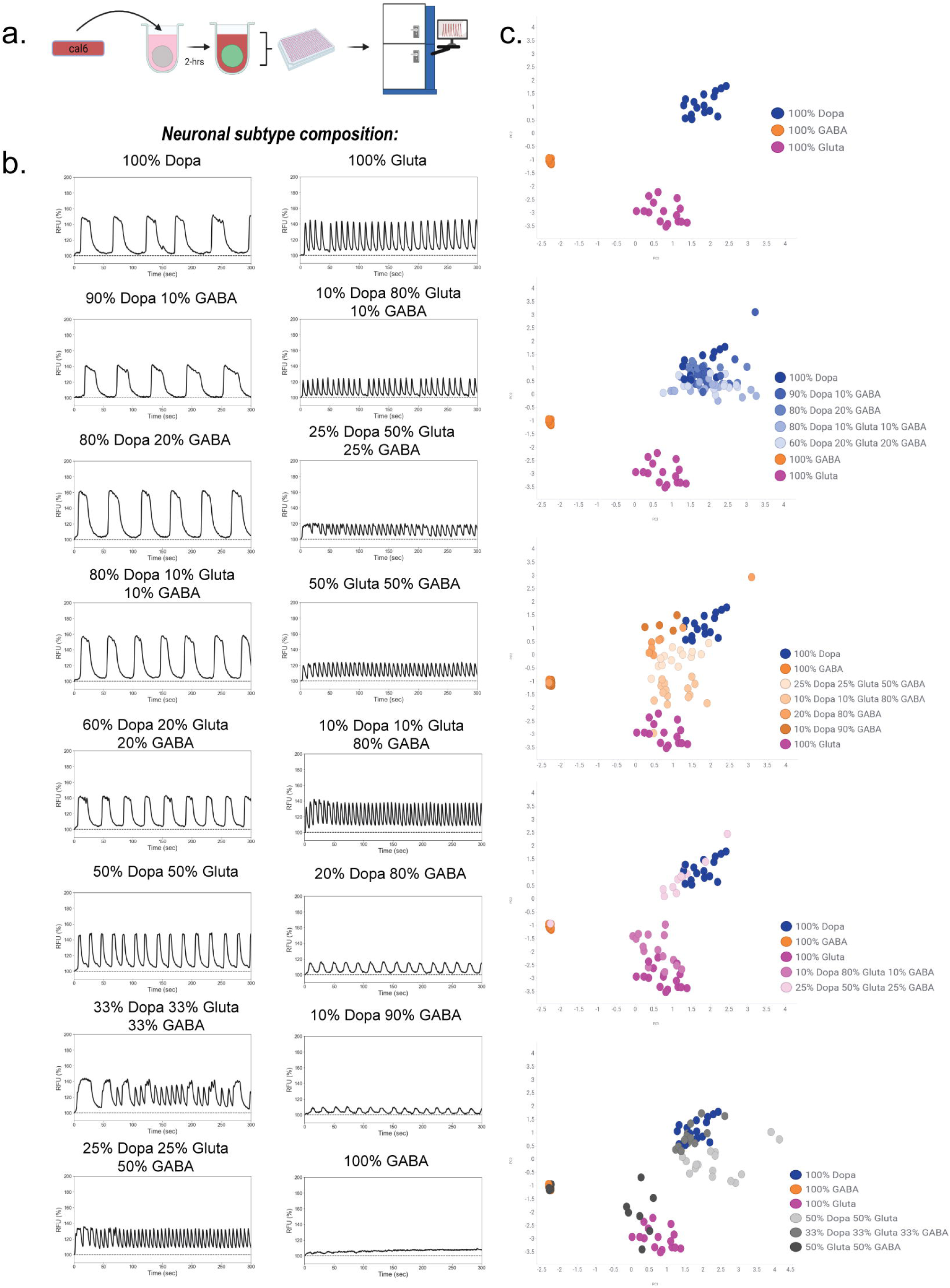
Calcium activity phenotypes are dependent on neuronal cell type composition. **(a.)** Schematic of experimental design; spheroids were incubated with Cal6 dye for 2-hrs prior to FLIPR recordings after a 3-week maintenance period. **(b.)** Representative traces from 16 different spheroids that are 90% neuron and 10% astrocyte but differ by neuronal subtype composition. **(c.)** Principal component analysis (PCA) was used as a dimension reduction algorithm to incorporate 10 peak parameters extracted from the multiparametric peak analysis for all wells, and scatter plots were used to visualize the spatial distribution of calcium activity phenotypes for each spheroid type. Plots from top to bottom: 1. single neuron spheroid (SNS) types (100% dopaminergic, glutamatergic, or GABAergic neurons) 2. Spheroids with majority dopaminergic neurons (shades of blue) relative to SNSs 3. spheroids with mostly GABAergic neurons (shades of orange) relative to SNSs 4. spheroids with mostly glutamatergic neurons (shades of pink) relative to SNSs 5. spheroids with equal distributions of neuronal cell types. Data was obtained from 16 technical replicates per spheroid type over one experiment; Raw data obtained from the multiparametric peak analysis was used for the PCA.

#### Calcium Activity Analysis

Calcium oscillatory peak detection data from FLIPR recordings was obtained through ScreenWorks 5.1 (Molecular Devices). Initial peak detection analysis occurred within ScreenWorks 5.1 via the PeakPro 2.0 module. Here, all parameters were set to be the same for all wells per plate. The event polarity was always set to positive and search vector length always set to 11. The baseline, trigger level, which is automatically set to 10% above the baseline, and dynamic threshold, which is the threshold for peak detection were automatically identified by the PeakPro 2.0 module. Wells were manually checked to ensure these parameters were accurately identified prior to analysis. After analysis, data from 18 peak parameters (mean peak amplitude, peak amplitude standard deviation (SD), peak count, mean peak rate, peak rate SD, peak spacing, peaking spacing SD, mean number of EAD-like peaks per well, CTD @ 50% (peak width at 50% amplitude), CTD @ 90% (peak width at 90% amplitude), rise slope, rise slope SD, mean peak rise time, peak rise time SD, decay slope, decay slope SD, mean peak decay time, and peak decay time SD) was exported to a STATALL file that could be converted to a Microsoft Excel spreadsheet. Percent coefficient of variance (%CV) was calculated within each plate similar to what is described in Boutin et al., 2022 to measure how variable each parameter exported was. Parameters were included in future analysis if they were under the threshold cutoff of 30% CV, which included peak count, peak rate, peak spacing, peak width 50% and 90%, peak amplitude, peak rise time, peak decay time, rise slope, and decay slope (Table S1). All data was normalized to the average of DMSO-treated control wells within each plate. Each group represented on radar plots shows the mean in comparison to DMSO vehicle controls, which should always average to 100%. Bar plots with individual values are reported as mean ± SEM.

For peak detection data obtained from the Phenix Plus confocal microscope, image sequences were stored in and exported from Columbus Image Data Storage and Analysis as single plane TIFFs. Each recording was imported into ImageJ and converted to a single stack. Prior to peak detection analysis, the T-function, F div F0, in ImageJ was used to obtain calcium signals normalized to background fluorescence. The ImageJ Plugin, LC_Pro, which was first described by Francis et al., 2014, was used to automatically identify regions of interest containing dynamic calcium signals across the image sequence. The automated analysis was used on the F div F0 recording so that calcium measurements would be reported as normalized fluorescent values (F/F0). For the LC_Pro analysis, default settings were used. F/F0 values for a region of interest (ROI) were exported if they contained a high signal to noise ratio and exported to a text file titled ROI Normalized. Once this text file was converted to a csv file, it was uploaded into Python for peak detection analysis. While code for this analysis is publicly available on GitHub, a brief description of the process is described. First, data from all identified ROIs was normalized such that the minimum F/F0 value was equal to 1. Time series plots showing mean signal plus variability represented as 95% confidence interval were generated, along with heatmaps showing activity across every identified ROI, and a correlation matrix plotted as a heatmap representing correlation coefficients across all ROIs. The correlation matrix was used to describe how synchronous the calcium activity within a spheroid was, and a synchrony score was measured by calculating the average correlation coefficient across the matrix. Given that LC_Pro identifies different numbers of ROIs for each recording, we used a random sample generator to randomly choose 12 ROIs for peak detection analysis. To ensure this random sample reflected the population activity, we required the correlation score of the random sample to be within 5% of the correlation score of the population of ROIs. The find_peaks package was imported from scipy.signal and used for peak detection analysis. The scipy package, find_peaks was used to detect and measure peak parameters including peak count, amplitude, and width (Virtanen et al., 2020).

#### Statistical Analysis

Python, R Studio and GraphPad Prism were used for statistical analysis. Principal component analysis was performed in Python using the package, scikit-learn (Pedregosa et al., 2011), and number of components was determined based on retaining > 85% of variance. Prior to quantifying predictive accuracy of activity phenotypes in neurological disease models with a random forest classifier (RFC) machine learning model, training and test datasets were generated which contained 80% and 20% of the data, respectively. A t-test was used to ensure no significant differences between data in the training versus the test sets, and z-tests were used to ensure similar frequencies of each genotype or pre-treatment group were represented. After confirming no significant differences between training and test datasets, the PCA data in the test dataset was tested against the RFC model and labeling accuracy for each genotype or pre-treatment group are reported. Comparisons between two groups were analyzed with unpaired t-tests when comparing separate groups of spheroids, and paired t-tests when examining effects within the same spheroid. Comparisons between two-groups or more were analyzed with one way ANOVA, and significant main effects were followed up with Dunnett’s multiple comparisons tests. Data is reported as mean ± SEM for column and violin plots, and as mean for radar plots. Significance was set at p < 0.05.

#### Reproducibility

Data used in the current study spanned ten 384-well plates. Data from Figure 2 consisted of 16 technical replicates per group from one independent experiment. Data from Figures 3-7 was run over three independent experiments, with technical replicates consisting of n > 3 per experiment. Data from Figure 8 was collected from two independent experiments with n > 2 technical replicates per experiment.

**Figure 3.**
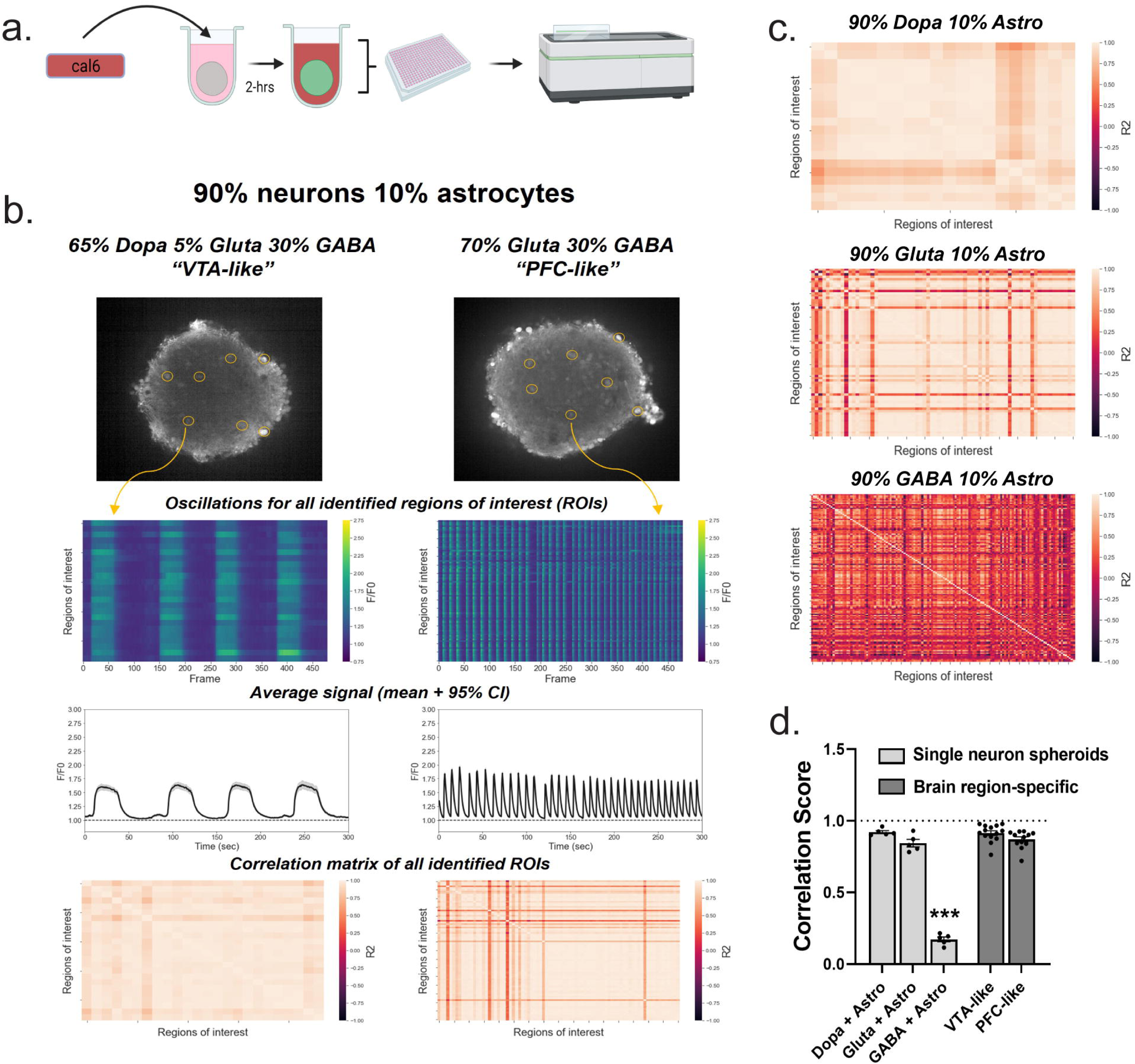
Synchronous calcium oscillations occur in brain region-specific spheroids as well as single neuron spheroids with dopaminergic and glutamatergic, but not GABAergic, neurons. **(a.)** Schematic of experimental design; spheroids were incubated in Cal6 dye for 2-hrs prior to confocal time series recordings after a 3-week maintenance period. **(b.)** Calcium activity in brain region-specific spheroids modeling the ventral tegmental area (VTA-like) and prefrontal cortex (PFC-like); Top: visual of the automated region of interest (ROI) detection used to examine calcium activity within several cells in each spheroid; Second from top: calcium activity from all identified ROIs shown in the form of a heatmap, with tick marks for ROIs in increments of 5 on the left y-axis, dF/F0 on the right y-axis, and video frame on the x-axis; Second from bottom: Average signal of all identified ROIs plus 95% confidence interval; Bottom: Correlation matrices displaying R2 values as a heatmap for all identified ROIs within a spheroid; Regions of interest (ROIs) with oscillatory patterns were automatically identified using the LC_Pro plugin through ImageJ and activity of all identified ROIs was plotted as a heatmap and population activity was represented by the mean plus 95% confidence interval of all identified ROIs (time series trace) **(c.)** Correlation matrices from representative single neuron spheroids showing synchrony of all identified ROIs **(d.)** Quantification of synchrony through a correlation score, obtained through the mean R2 value for each spheroids’ correlation matrix. Data was run in technical replicates of n=4-6 per plate across one experiment for SNSs and three separate experiments for brain region specific spheroids; For d., One way ANOVA (F_(4,35)=_100.7, p<0.0001) was followed up with Dunnett’s posthoc (p<0.0001 for all groups compared to GABA + Astro spheroids), data is represented as mean ± SEM, *** p < 0.001.

## RESULTS

### Designer neural spheroids exhibit differential calcium activity profiles depending on neuronal subtype composition

We first sought to establish whether iPSC-derived, differentiated neurons could be incorporated into a co-culture spheroid system and have functional neuronal activity. We mixed glutamatergic GABAergic, and dopaminergic neurons with astrocytes and seeded as cell mixtures of controlled ratios into 384-well, ultra-low attachment, round bottom plates to force cell aggregation into the formation of spheroids (**Fig. 1a**). We observed spheroid formation after 3 days in culture in both PFC-like spheroids (70% glutamatergic, 30% GABAergic neurons) and VTA-like spheroids (65% dopaminergic, 5% glutamatergic, 30% GABAergic neurons), both consisting of 90% neurons and 10% astrocytes. The neuronal composition of the “VTA-like” and “PFC-like” spheroids was based on the cell-type composition of these regions from postmortem human brain studies (Lin et al., 2013; Pignatelli et al., 2015; Root et al., 2016). Calcium activity was recorded after 21 days using a calcium fluorescence (Cal6) dye with a whole plate reader equipped with a high speed, high sensitivity EMCCD camera for both fluorescent detection (the FLIPR Penta High-Throughput Cellular Screening System, referred from here on as “FLIPR”), suggesting formation of active, synchronized neuronal activity. Spheroids formed by this protocol were <400 μm in diameter (**Fig 1b**) after the maturation process and had a homogenous spatial distribution of neurons and astrocytes (**Fig 1b**). Furthermore, immunostaining for tyrosine hydroxylase (TH) showed that a majority of neurons in VTA-like spheroids were dopaminergic while only a few expressed TH in PFC-like spheroids (**Fig 1c, Video S1**). Parvalbumin (PV) staining revealed similar compositions of GABAergic neurons between PFC- and VTA-like spheroids, which were each generated with 30% GABAergic neurons (**Fig 1c, Video S2**). We were able to detect higher signal of vGluT1 protein expression in PFC-like spheroids (**Fig. 1c, Video S2**), in addition to higher mRNA expression of vGluT1 with RNAscope **(Fig. S1**). The mature functional spheroids also expressed pre- and postsynaptic markers as shown by synapsin and homer staining, respectively, distributed evenly throughout spheroids, suggesting the presence of synaptic connections (**Fig 1d**).

To test whether this spheroid system was robust for high-throughput (HT) screening, we measured calcium activity across all wells per plate simultaneously using a FLIPR. We then analyzed the measured calcium oscillations from spheroids incubating in Cal6 dye for high reproducibility peak parameters using ScreenWorks PeakPro 2.0 analysis (**Fig 2a**). Specifically, we analyzed 17 peak parameters and selected 10 reproducible parameters with low variability (<30% coefficient of variance (%CV), (**Table S1**). We performed a proof-of-concept study measuring calcium activity in 16 different spheroid types that all contained 90% neurons and 10% astrocytes but differed in their neuronal subtype composition to assess whether changing neuronal cell type composition would impact phenotypic profiles (**Fig. 2b**). Principal component analysis (PCA) was used to analyze the multiparametric peak data (**Fig 2c**). We observed that single neuron spheroids (SNSs, 90% neuron 10% astrocyte) consisting of only dopaminergic, glutamatergic, or GABAergic neurons form distinct clusters (**Fig. 2c, topmost graph**), suggesting cell-type specific phenotypic profiles, and that spheroids with controlled gradient ratios of multiple neuronal cell types cluster near the SNS cluster with the dominant neuronal cell type (**Fig. 2c**,). Together, this data shows that altering neuronal cell type composition within spheroids produces unique calcium activity phenotypes that reflect neuronal subtype ratios.

After establishing culture maturation conditions and maintenance of individual and heterogenous neuronal subtypes, we next characterized PFC- and VTA-like spheroids, described in **Fig. 1**, by confocal microscopy to measure calcium activity from individual cells within either the single neuron spheroids (SNSs) or brain region-specific spheroids (schematic, **Fig 3a, Video S3, S4)**. In contrast to the FLIPR assay that measures population activity, synchrony between cells could be measured with confocal microscopy. Correlation matrices generated from individual cell activity between all identified regions of interest (ROIs) were plotted as a heatmap, and the average correlation coefficient (R^2^) value from each spheroid was calculated as its “correlation score” (**Fig 3b,c**). VTA- and PFC-like spheroids displayed unique phenotypic profiles similar to the phenotypes displayed by SNSs with dominant neuronal cell types (**Fig 3b, S2a)**. In addition, synchronous activity was observed in SNSs with dopaminergic and glutamatergic neurons along with both brain region-specific spheroid types, but not in SNSs with GABAergic neurons (**Fig 3c,d**). While we observed that astrocytes were not necessary for activity, astrocyte presence altered peak parameters in SNSs containing dopaminergic or glutamatergic neurons. Given the role astrocytes play in neurotransmitter release, synaptic plasticity and neurological diseases (Chung et al., 2015; Ota et al., 2013; Perez-Catalan et al., 2021), spheroids tested throughout the rest of the study were all made with 90% neurons and 10% astrocytes **(Fig. S2a, 2b)**. Together, these data indicate that brain region-specific spheroids display unique phenotypic profiles, and that changes in synchronous neuronal activity in these spheroids occurs through GABAergic mechanisms.

### Phenotypic profiles in brain region-specific spheroids can be differentially modulated with compounds targeting neuronal subtype receptors

We next validated spheroid functional response via treatment with quality control (QC) compounds that targeted receptors on each neuronal subtype (**Fig. 4**, **Fig. S3, Table S2)**. In both spheroid types, treatment with the GABA_A_ receptor (GABA_A_R) agonist muscimol led to complete inhibition of activity **(Fig. 4, S3, Table S2)**. Conversely, treatment with GABA_A_R antagonist, bicuculline, induced spheroid type-dependent changes; in VTA-like spheroids, enhanced activity was apparent through increases in peak count, rate, and amplitude with decreases in peak rise and decay time while in PFC-like spheroids, peak count was reduced while peak amplitude and peak width was increased, suggesting different functional responses between the spheroids to GABA_A_R antagonism (**Fig. 4**, **Table S2)**. To target glutamatergic neurons, spheroids were treated with the α-amino-hydroxy-5-methyl-4-isoxazolepropionic acid receptor (AMPAR) antagonist, CNQX, which inhibited all activity in PFC-like spheroids but led to increases in peak rise time and peak count in VTA-like spheroids (**Fig. 4**, **Table S2).** The N-methyl-D-aspartate receptor (NMDAR) antagonist, memantine. Memantine increased peak count and rate and decreased amplitude in both spheroid types (**Fig. 4**, **Table S2)**. SCH23390 was used to block stimulatory G-protein coupled dopamine 1/5 receptors (D1Rs) while sulpiride was used to block inhibitory G-protein coupled dopamine 2/3 receptors (D2Rs). In both spheroid types, D1R antagonism with CNQX led to total inhibition in both spheroid types while D2R antagonism with sulpiride increased peak count in PFC-like spheroids and peak width in VTA-like spheroids (**Fig. 4**, **Table S2)**. Together, this data shows that predictable functional responses occurred in brain region-specific spheroids based on their neuronal cell type composition.

**Figure 4.**
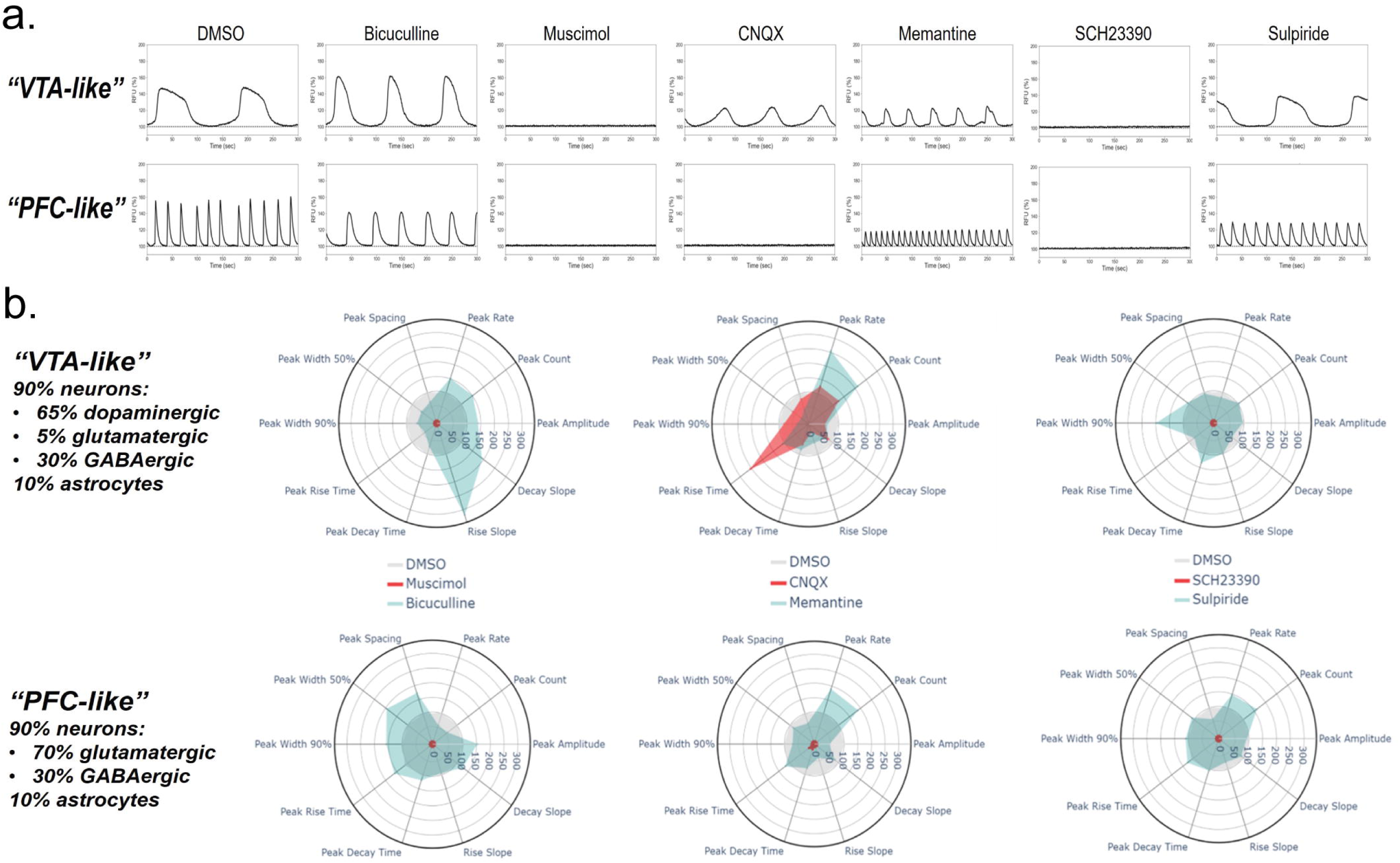
Functional responses to quality control (QC) compounds in brain region-specific neural spheroids. **(a,b.)** Data collected from FLIPR recordings obtained from spheroids in a Cal6 dye **(a.)** Representative time series plots showing calcium activity phenotypes after treatment with control compounds targeting receptors for each neuronal cell type from VTA-like spheroids on top and PFC-like spheroids on the bottom. Left to right: DMSO (vehicle), Bicuculline (GABA_A_R antagonist), Muscimol (GABA_A_R agonist), CNQX (AMPAR antagonist), Memantine (NMDAR antagonist), SCH23390 (Dopamine 1 receptor (D1R) antagonist), and Sulpiride (D2R antagonist). **(b.)** Radar plots depicting multiparametric calcium activity response to each compound relative to DMSO controls in VTA-like (top) and PFC-like (bottom) spheroids. For each compound, wells were tested at n=3-4 technical replicates over three separate experiments; For b., results are normalized to DMSO-treated control wells such that averages for DMSO-treated control wells on the radar plot equal 100%. Data from b. are represented as mean values for each parameter.

### Incorporating genetically engineered GABA neurons expressing APOE e4/4 allele produces a predictive calcium activity phenotype that is reversed with clinically approved treatments for Alzheimer’s Disease

To model Alzheimer’s Disease (AD), we used GABA neurons that were genetically engineered to carry the apolipoprotein e4/4 (APOE4) allele, a genotype associated with AD (Belloy et al., 2019; Lin et al., 2018). PFC-like spheroids were made either containing 30% APOE3 (Wt) or APOE4 (mutant) GABA neurons. SNSs with Wt or APOE4 GABA neurons were made as controls. After a 3-week maturation period, there was no significant differences in spheroid viability between APOE3 and APOE4 GABA neurons, suggesting that functional differences observed between genotypes would not be due to cell viability differences caused by the APOE4 mutation **(Fig. S4a)**.

Both PFC-like and single neuron GABA spheroids with APOE4 GABA neurons displayed reduced peak count by ROI analysis (**Fig. 5a,b**). However, peak amplitude was reduced only in SNSs with APOE4 GABA neurons while peak width was decreased in SNSs but increased in PFC-like spheroids with APOE4 GABA neurons (**Fig. 5a,b**). In PFC-like spheroids, the incorporation of APOE4 GABA neurons also disrupted synchronous neuronal activity, as indicated by reduction in correlation scores (**Fig. 5a,b**). FLIPR recordings of baseline activity replicated the confocal recording, showing that APOE4 GABA neurons in PFC-like spheroids caused a reduction in peak count and an increase in peak width (**Fig. 5c**). Given the non-synchronous activity of SNSs with GABAergic neurons, the FLIPR was unable to detect peaks in these spheroids so genotypic differences were not observed (**Fig. 5c**). We implemented a random forest classifer (RFC) supervised machine learning algorithm, described in methods, to measure labeling accuracy of genotypes based on baseline FLIPR data in PFC-like spheroids to quantify how predictive the APOE4 phenotype was for the AD model. The RFC model accurately predicted the label of 96% of Wt (APOE3) PFC-like spheroids and 92% of APOE4 PFC-like spheroids (**Fig. 5e**). These data show that APOE4 GABA neurons produce baseline phenotypic deficits that are highly predictive when tested against the RFC machine learning algorithm.

**Figure 5.**
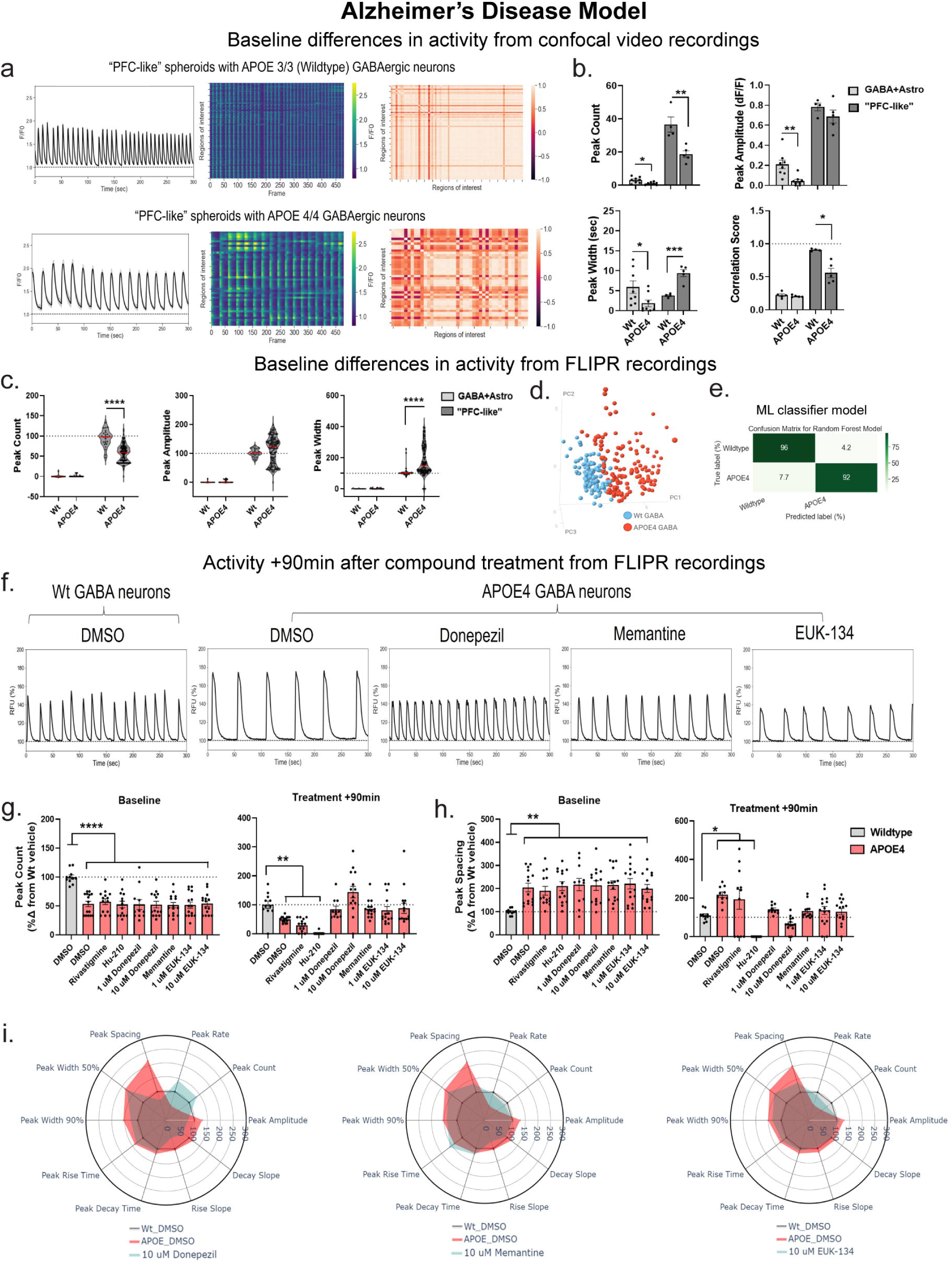
Clinically approved compounds to treat symptoms of Alzheimer’s Disease (AD) reverse deficits caused by the incorporation of APOE4/4 GABA neurons in PFC-like spheroids to model AD. **(a,b.)** Data collected from confocal recordings **(a.)** Baseline phenotypic differences in PFC-like spheroids containing GABA neurons expressing APOE3/3 (Wildtype, top panel) or APOE4/4 (bottom panel) to model AD; Left: average signal across all identified ROIs, Middle: heatmap showing activity of all identified ROIs, Right: correlation matrix showing synchrony of ROIs **(b.)** Quantification of peak count (SNS: t_(14)_=2.92, p=0.011; PFC-like: t_(7)_=3.81, p=0.007), amplitude (SNS: t_(14)_=3.58, p=0.011), width (SNS: t_(14)_=2.27, p=0.039; PFC-like: t_(7)_=5.46, p=0.0009), and synchrony from confocal recordings (PFC-like: t_(7)_=4.36, p=0.003) (**c-i.)** Data collected from FLIPR recordings **(c.)** Baseline peak count (SNS: p=0.25; PFC-like: p<0.0001), amplitude (SNS: p=0.21; PFC-like: 0.056), and width (SNS: p=0.18; PFC-like: p<0.0001) from all wells recorded with FLIPR **(d.)** PCA on baseline multiparametric peak data represented as a scatterplot displaying individual values **(e.)** Predictive labeling of genotype based on PCA data is 94% accurate using Random Forest machine learning classifier model; Data represented as a confusion matrix showing accurate and erroneous error labels for each genotype, values represented as percentages **(f.)** Representative time series traces 90min after Wt and APOE4 spheroids were treated with compounds used to treat AD **(g,h.)** Peak count **(g.)** and peak spacing **(h.)** in Wt PFC-like spheroids treated with DMSO and APOE4 spheroids divided by treatment group at baseline (left) and 90min after treatment (right); Donepezil, Memantine, and EUK-134 restore deficits in peak count (peak count, baseline: F_(8,115)_=7.05; p<0.0001; treatment: F_(8,114)_=12.4, p<0.0001; peak spacing, baseline: F_(8,115)_=3.06, p=0.004; treatment: F_(8,102)_=10.22, p<0.0001) **(i.)** Radar plots showing phenotypic data across 10 peak parameters measured 90min after treatment with DMSO in Wt (gray) and APOE4 (red) spheroids, along with treatment with either Donepezil, Memantine, and EUK-134 (teal). For confocal recordings, n=4-8 per group over 2 separate experiments; For FLIPR recordings, n= 249 samples over 3 separate experiments. Data from (b,g,h.) represented as mean ± SEM, (b.) analyzed with unpaired t-tests for each spheroid type & (g,h.) analyzed with One way ANOVA followed up with Dunnett’s posthoc. Data from (c.) represented as violin plots with median indicated by red line; Mann-Whitney unpaired t-test used to compare medians between groups, (i.) represented as radar plots showing group averages for each peak parameter analyzed. * p < 0.05, **p<0.01, ***p<0.001, ****p<0.0001).

We next sought to find out whether APOE4-induced deficits in PFC-like spheroids could be reversed following treatment with three clinically approved compounds used to treat the symptoms of AD in humans along with two preclinical compounds that are known to inhibit beta-amyloid plaques. To do this, FLIPR recordings were obtained 90-min after compound treatment in the same spheroids analyzed in **Fig 5c-e**. The clinically approved compounds included cholinesterase inhibitors (Rivastigmine and Donepezil) along with an NMDAR antagonist (Memantine) while the preclinical compounds were Hu-210 and EUK-134, which have both been shown to inhibit beta-amyloid plaque production in animal models (Bahramikia and Yazdanparast, 2013; Chen er et al., 2010; Jekabsone et al., 2006; Ramirez et al., 2005). All data was normalized to Wt DMSO-treated controls. Baseline deficits were consistent between all treatment groups prior to compound addition, and show significant reductions in peak count and increases in peak spacing among all treatment groups in APOE4 PFC-like spheroids compared to Wt controls (**Fig. 5g,h**). Three compounds were found to reverse deficits caused by APOE4 GABA neurons including Memantine (10 µM), Donepezil (1 & 10 µM), and EUK-134 (1 & 10 µM) since deficits in peak count and spacing were no longer significantly different from Wt DMSO-treated spheroids (**Fig. 5f-h**, panels labeled Treatment +90min). In Wt PFC-like spheroids, these same compounds also increased peak count and decreased peak spacing, which may indicate a general rather than APOE4 specific mechanism **(Fig. S5)**. Radar plots for each of these three compounds illustrate the phenotypic profiles across all peak parameters 90min after treatment compared to both DMSO-treated controls (**Fig 5i**). These findings show that functional deficits caused by the APOE4 mutation in GABA neurons can be partially reversed by compounds used to treat AD symptoms and lays the groundwork for HTS studies using spheroid models of AD.

### An A53T SNCA model of Parkinson’s Disease produces predictive phenotypic deficits in calcium activity that can be reversed with a dopamine agonist

We next modeled Parkinson’s Disease (PD) by incorporating dopaminergic neurons expressing mutant A53T alpha-synuclein into spheroids given that it is a common risk factor for non-familial PD (Fernandes et al., 2020; Petrucci et al., 2016; Zambon et al., 2019). A53T dopaminergic neurons were used to make mutant VTA-like spheroids along with Wt or A53T dopaminergic SNSs consisting of 90% neurons and 10% astrocytes. VTA-like spheroids were formed with 65% Wt or A53T dopaminergic, 5% glutamatergic, and 30% GABAergic neurons. No differences in spheroid viability between Wt and A53T neurons for either spheroid type were observed **(Fig. S4b)**.

Confocal recording revealed increased peak count and decreased peak amplitude and width in both VTA-like and SNSs with A53T dopaminergic neurons, but similar levels of synchrony (**Fig. 6a,b**). We also measured whole spheroid activity with the FLIPR and similarly saw that A53T dopaminergic neurons increased peak count and decreased peak amplitude and width in both spheroid types (**Fig. 6c**). We found a significant increase in subpeak count, amplitude, and width in VTA-like spheroids with A53T neurons but not in SNSs with A53T dopaminergic neurons (**Fig. 6d**). The RFC model tested on baseline FLIPR data found that 97% of Wt and 95% of A53T VTA-like spheroids were accurately labeled, giving an average accuracy score of 96% (**Fig. 6f**). Together, this suggests that the baseline deficits induced by the incorporation of A53T dopaminergic neurons in VTA-like spheroids creates a highly predictive, and highly reproducible, calcium activity phenotype.

**Figure 6.**
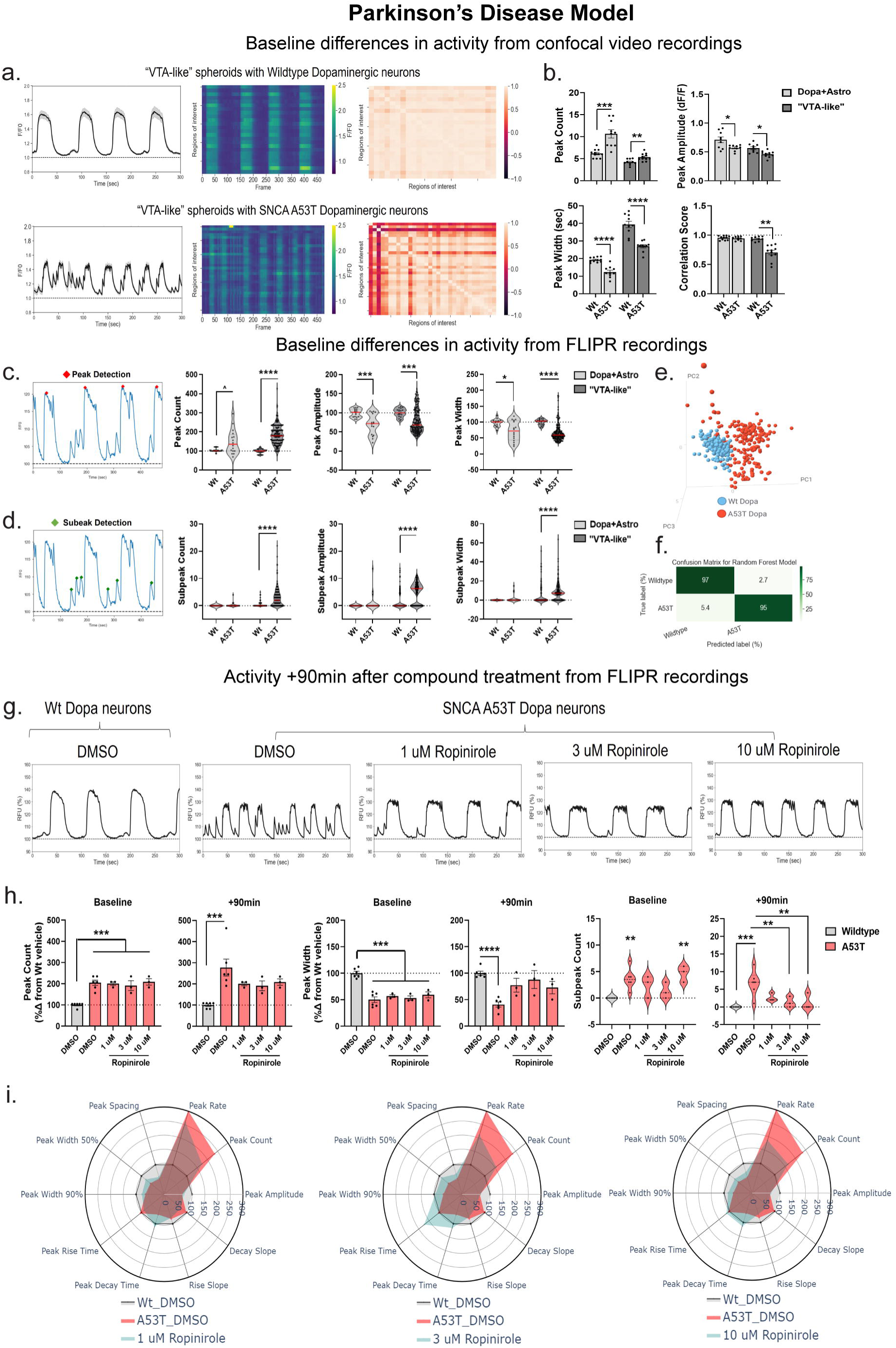
Dopamine agonist, Ropinirole, reverses deficits induced by incorporation of mutant alpha-synuclein (SNCA A53T) dopaminergic neurons into VTA-like spheroids to model Parkinson’s Disease (PD). **(a,b.)** Data collected from confocal recordings **(a.)** Baseline phenotypic differences in VTA-like spheroids containing wildtype (top panel) or mutant A53T dopaminergic neurons (bottom panel) to model PD; Left: average signal across all identified ROIs, Middle: heatmap showing activity of all identified ROIs, Right: correlation matrix showing synchrony of ROIs **(b.)** Quantification of peak count (SNS: t_(19)_=4.73, p =0.0001; VTA-like: t_(19)_=3.37, p=0.0032), amplitude (SNS: t_(16)_=2.71, p =0.015; VTA-like: t_(18)_=3.66, p=0.002), width (SNS: t_(19)_=6.55, p<0.0001; VTA-like: t_(18)_=6.09, p<0.0001), and synchrony (VTA-like: t_(18)_=5.29, p<0.0001) from confocal recordings **(c-i.)** Data collected from FLIPR recordings **(c.)** Example trace showing peak detection analysis as well as quantification of baseline peak count (SNS: p=0.055; VTA-like: p<0.0001), amplitude (SNS: p=0.0007; VTA-like: p=0.0002), and width (SNS: p=0.027; VTA-like: p<0.0001) from all wells recorded with FLIPR **(d.)** Example trace of VTA-like A53T spheroid showing subpeak detection analysis as well as quantification of baseline subpeak count (SNS: p=0.23; VTA-like: p<0.0001), amplitude (SNS: p=0.23; VTA-like: p<0.0001), and width (SNS: p=0.23; VTA-like: p<0.0001) from all wells recorded with FLIPR **(e.)** PCA on baseline multiparametric peak data represented as a scatterplot displaying individual values **(f.)** Predictive labeling of genotype based on PCA data is 96% accurate using Random Forest machine learning classifier model; Data represented as a confusion matrix showing accurate and erroneous error labels for each genotype, values represented as percentages **(g.)** Representative time series traces 90min after Wt and A53T spheroids were treated with DMSO or Ropinirole, a dopamine agonist **(h.)** Peak count (baseline: F_(4,16)_=17.6, p<0.0001; treatment: F_(4,16)_=7.5, p=0.0001), peak width (baseline: F_(4,16)_=23.27, p<0.0001; treatment: F_(4,16)_=8.99, p<0.0001), and subpeak count (baseline: F_(4,16)_=6.69, p=0.002; treatment: F_(4,16)_=7.17, p=0.0017) in Wt VTA-like spheroids treated with DMSO and A53T spheroids treated with Ropinirole, which restored deficits across all 3 peak measures **(i.)** Radar plots showing phenotypic data across 10 peak parameters measured 90min after treatment with DMSO in Wt (gray) and A53T (red) spheroids, along with treatment with Ropinirole across 3 concentrations (teal). (For confocal recordings, n=8-10 per group over 2 separate experiments; For FLIPR recordings, n= 370 samples over 3 separate experiments. Data from (b,h.) represented as mean ± SEM, (b.) analyzed with unpaired t-tests for each spheroid type & (h.) analyzed with One way ANOVA followed up with Dunnett’s posthoc. Data from (c.) represented as violin plots with median indicated by red line; Mann-Whitney unpaired t-test used to compare medians between groups, (i.) represented as radar plots showing group averages for each peak parameter analyzed. * p < 0.05, **p<0.01, ***p<0.001, ****p<0.0001).

After the baseline FLIPR recording, spheroids were treated with clinically approved treatments for PD to measure whether they could reverse disease-related deficits in VTA-like spheroids with A53T dopaminergic neurons. These treatments included L-Dopa, used as dopamine replacement therapy, Ropinirole (dopamine agonist), Entacapone and Tolcapone (catechol-O-methyltransferase (COMT) inhibitors), Rasagiline (monoamine oxidase type B (MAO-B) inhibitor), Benztropine (dopamine transporter inhibitor), Trihexyphenidyl (antimuscarinic), and Amantadine (antiviral). Treatment effects were measured with FLIPR recordings 90-min after treatment. Only Ropinirole (1,3,10 µM) reversed peak count and increased peak width 90-min after treatment to the level that they were no longer significantly different from Wt DMSO-treated control spheroids (**Fig. 6h, S5**). Furthermore, 3 and 10 µM Ropinirole significantly reduced subpeak count to where they were similar to Wt DMSO-treated controls (**Fig. 6h**). Differences across all peak parameters are displayed as radar plots, where phenotypic profiles for all three concentrations of Ropinirole were tested (**Fig. 6i**). In summary, the dopamine agonist Ropinirole was able to reverse deficits induced by the incorporate of A53T mutant dopaminergic neurons, showing that our PD model responds to treatments clinically approved to treat PD in humans.

### Naloxone reverses deficits induced by chronic DAMGO treatment in PFC-like but not VTA-like spheroids

We next modeled opioid use disorder (OUD) by developing a protocol intended to model various facets of addiction including drug intake and withdrawal given the current opioid epidemic (Rehm and Shield, 2019). We modeled drug intake by chronically treating spheroids with 10 µM DAMGO, a µ-opioid receptor agonist, in the media every other day for 10-days. DAMGO withdrawal was modeled in the same way, but spheroids were not treated on the final day such that this treatment group received control media 3-days prior to recording (**Fig. 7a**). DAMGO pre-treatment did not impact spheroid viability (**Fig. S4c**).

**Figure 7.**
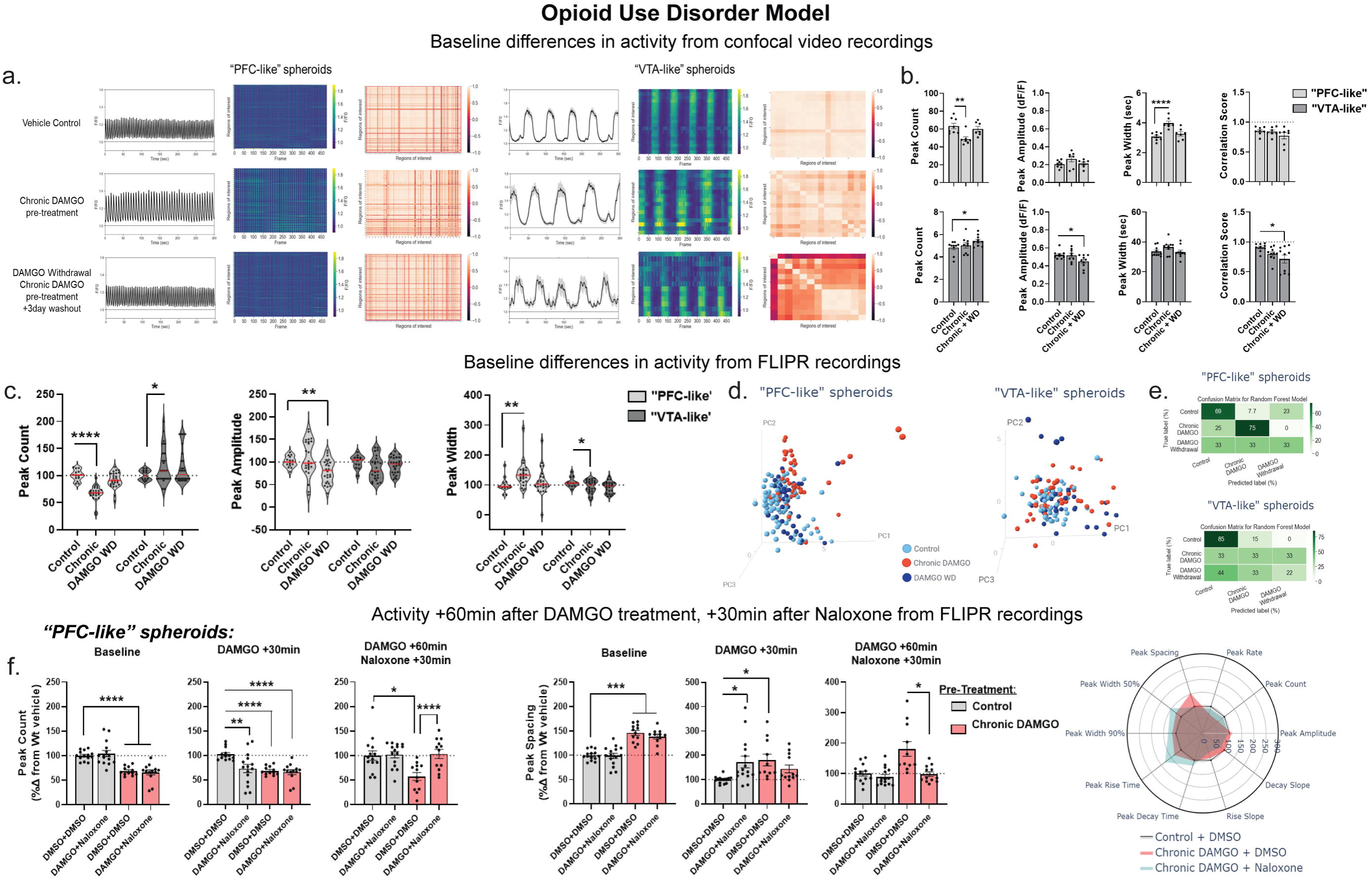
Naloxone (MOR antagonist) reverses deficits induced by chronic opioid treatment in PFC-like but not VTA-like spheroids in spheroids modeling Opioid Use Disorder. **(a,b.)** Data collected from confocal recordings **(a.)** Baseline phenotypic differences in PFC- and VTA-like spheroids pre-treated with either vehicle (water, top panel), chronic DAMGO (middle panel), or chronic DAMGO plus a 3-day washout to model drug withdrawal; Left: average signal across all identified ROIs, Middle: heatmap showing activity of all identified ROIs, Right: correlation matrix showing synchrony of ROIs **(b.)** Quantification of peak count (PFC-like: F_(2,19)_=7.17, p=0.005; VTA-like: F_(2,26)_=2.26, p=0.097), amplitude (PFC-like: F_(2,19)_=1.99, p=0.16; VTA-like: F_(2,26)_=4.03, p=0.03), width (PFC-like: F_(2,19)_=14.91, p=0.0001; VTA-like: F_(2,26)_=2.04, p=0.15), and synchrony (PFC-like: F_(2,19)_=0.74, p=0.28; VTA-like: F_(2,26)_=4.27, p=0.025) from confocal recordings **(c-f.)** Data collected from FLIPR recordings **(c.)** Quantification of baseline peak count, amplitude, and width from all wells recorded with FLIPR (peak count: PFC-like: F_(2,61.7)_=36.9, p<0.0001, VTA-like: F_(2,53.5)_=2.04, p=0.14; amplitude: PFC-like: F_(2,47.18)_=7.23, p=0.002, VTA-like: F_(2,54.4)_=2, p=0.15; width: PFC-like: F_(2,40.3)_=5.36, p=0.007 VTA-like: F_(2,53.5)_=3.33, p=0.04) **(d.)** PCA on baseline multiparametric peak data represented as a scatterplot displaying individual values in PFC-like (left) and VTA-like spheroids (right) **(e.)** Predictive labeling of pre-treatment group based on PCA data using Random Forest machine learning classifier model; Data represented as a confusion matrix showing accurate and erroneous error labels for each pre-treatment group, values represented as percentages **(f.)** Peak count and peak spacing in PFC-like spheroids; Data shows baseline differences between spheroids with vehicle vs chronic DAMGO pre-treatment (left panel), 30min after DAMGO treatment (middle panel), and 60min after DAMGO treatment plus 30min after naloxone treatment; Data shows that naloxone is able to reverse deficits induced by chronic DAMGO pre-treatment and DAMGO treatment **(g.)** Radar plots showing phenotypic data across 10 peak parameters measured 60min after treatments with either DMSO, DAMGO, and/or naloxone. (For confocal recordings, n=7-8 per group over 2 separate experiments; For FLIPR recordings, n= 171 (PFC-like) and 135 samples (VTA-like) over 3 separate experiments. Data from (b,f.) represented as mean ± SEM, analyzed with One way ANOVA followed up with Dunnett’s posthoc. Data from (c.) represented as violin plots with median indicated by red line; Brown-Forsythe and Welch One way ANOVA used to compare medians between groups, (i.) represented as radar plots showing group averages for each peak parameter analyzed. * p < 0.05, **p<0.01, ***p<0.001, ****p<0.0001).

We observed by confocal microscopy chronic DAMGO treatment decreased peak count and increased peak width in PFC-like spheroids, while DAMGO washout spheroids modeling withdrawal did not have a significant change from control spheroids in peak count, amplitude, or width (**Fig. 7a,b**). In VTA-like spheroids, chronic DAMGO pre-treatment did not change baseline activity but DAMGO withdrawal significantly increased peak count, decreased peak amplitude, and disrupted synchrony compared to control spheroids (**Fig. 7a,b**). We observed similar differences using the FLIPR as were observed with the automated confocal recordings (**Fig. 7c**). The RFC model was preceded by PCA, and here we found 69% accurate labeling of phenotypes in chronic DAMGO PFC-like spheroids but only 33% accuracy labeling DAMGO withdrawal spheroids (**Fig. 7d,e**). For VTA-like spheroids, the RFC model accurately predicted 85% of control spheroids but only accurately identified 33% and 22% of chronic DAMGO-treated or DAMGO withdrawal spheroids, respectively (**Fig. 7d,e**), suggesting the phenotypic differences are not predictive in a classifier model.

Spheroids were then treated with 10 µM DAMGO, and calcium activity was recorded 30-min later, followed immediately by 10 µM naloxone, an MOR antagonist used to reverse opioid overdoses in humans. Both control and chronic DAMGO-treated spheroids were either treated with DMSO prior to each recording (DMSO+DMSO) or DAMGO followed by naloxone (DAMGO+Naloxone). 30-min after treatment with either DMSO or DAMGO, acute DAMGO significantly reduced peak count in control spheroids and chronic DAMGO pre-treated spheroids continued showing lower peak count (**Fig. 7f**, panels titled DAMGO +30min). For peak spacing, acute DAMGO treatment in control spheroids increased peak spacing, and peak spacing remained significantly increased in chronic DAMGO-treated spheroids that were treated with DMSO, but not DAMGO (**Fig. 7f**, panels titled DAMGO +30min). Spheroids were then treated with either DMSO or Naloxone and recorded 30-min later. We observed naloxone reversed deficits in peak count and spacing in both control spheroids acutely treated with DAMGO as well as chronic DAMGO pre-treated spheroids (**Fig. 7f**, panels titled DAMGO +30min, Naloxone +60min). These findings show that DAMGO treatment causes deficits in peak count and spacing that can be reversed by blocking µ-opioid receptors in PFC- but not VTA-like spheroids (**Fig. 7e**).

### Functional assembloids made from conjoining VTA- and PFC-like spheroids can be used to model neural circuitry

To extend this technology to model neural circuitry, we generated assembloids with the VTA and PFC spheroids. To activate the activity of neural circuits in assembloids, we first tested whether could be manipulate calcium activity using a chemogenetic approach (schematic shown in **Fig. 8a**). We transduced spheroids with retrograde AAV viruses inserting either stimulatory (hM3Dq) or inhibitory (hM4Di) designer receptors exclusively activated by designer drugs (DREADDs). Clozapine-N-oxide (CNO) was used at a 1 µM concentration to activate the DREADDs viruses, and representative time series traces as well as radar plots show the effect 30-min after treatment with FLIPR recordings (**Fig. 8b**). For dopaminergic and glutamatergic SNSs, chemogenetic activation increased peak count and rate while reducing parameters such as peak spacing and peak width, while the opposite was observed for these spheroids subjected to chemogenetic inhibition (**Fig. 8b,c**). We further showed that spheroids can express the genetically encoded calcium indicator, GCaMP6f, and that phenotypes within single VTA-like and PFC-like spheroids resemble what we observed from spheroids incubating in Cal6 dye (**Fig. S8a,b**).

**Figure 8.**
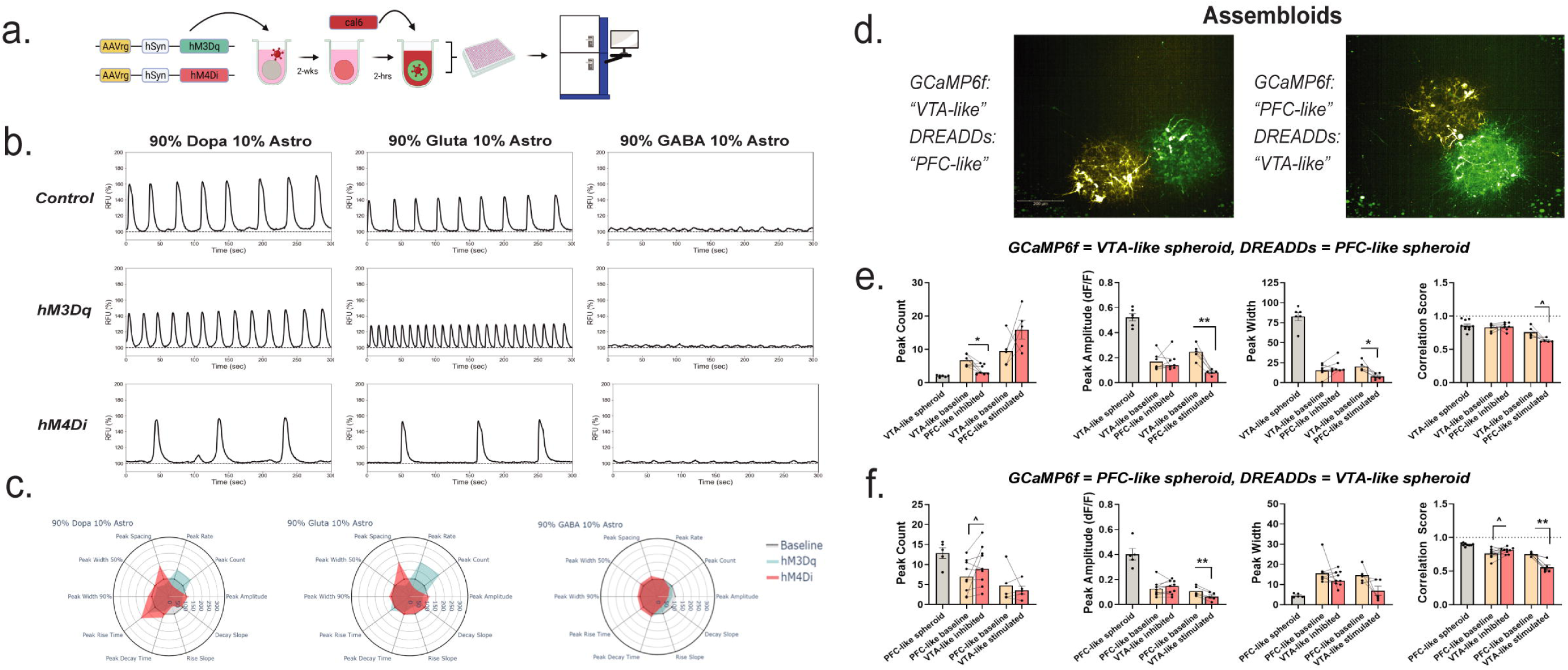
Functional assembloids can be made from conjoined spheroids to model neural circuitry. **(a.)** Schematic showing that spheroids can be transfected with DREADDs viruses tagged with mCherry and activity can be recorded with a FLIPR at 3-weeks using a cal6 dye. **(b.)** Activity from single neuron spheroids (SNSs) comprised of 90% neuron and 10% astrocytes can be recorded and displays similar phenotypes in spheroids transfected with stimulatory (hM3Dq) or inhibitory (hM4Di) DREADDs viruses. Top panel: vehicle control-treated spheroids expressing no DREADDs virus; Middle panel: 60-min after treatment with CNO to activate DREADDs virus and induce stimulatory activity; Bottom panel: 60-min after treatment with CNO to activate DREADDs virus and induce inhibitory activity **(c.)** Radar plots showing multiparametric peak alterations across 10 peak parameters for both stimulatory (teal) and inhibitory (red) DREADDs in SNSs with dopaminergic, glutamatergic, and GABAergic neurons, respectively. **(d.)** Representative image of two assembloids where one component expresses GCaMP6f and the other expresses either stimulatory (hM3Dq) or inhibitory (hM4Di) DREADDs viruses; Left: VTA-like expressing GCaMP6f, PFC-like expressing hM4Di, Right: PFC-like expressing GCaMP6f, VTA-like expressing hM4Di **(e,f.)** Quantification of baseline vs 90-min activity for peak count (hM4Di: t_(6)_=3.15, p=0.019; hM3Dq: t_(4)_=1.6, p=0.19), amplitude (hM4Di: t_(6)_=0.04, p=0.97; hM3Dq: t_(4)_=7.6, p=0.002), width (hM4Di: t_(6)_=1.93, p=0.1; hM3Dq: t_(4)_=3.29, p=0.03), and synchrony (hM4Di: t_(6)_=0.35, p=0.74; hM3Dq: t_(4)_=2.7, p=0.05) of the VTA-like component **(e.)** or PFC-like **(f.)** component (peak count: hM4Di: t_(8)_=1.94, p=0.09, hM3Dq: t_(4)_=0.65, p=0.55; amplitude: hM4Di: t_(8)_=0.8, p=0.45, hM3Dq: t_(4)_=7.3, p=0.002; width: hM4Di: t_(8)_=1.003, p=0.35, hM3Dq: t_(4)_=2.11, p=0.103; synchrony: hM4Di: t_(8)_=2.28, p=0.052, hM3Dq: t_(4)_=4.95, p=0.008) of assembloids before (yellow) and after (red) CNO was added to media to activate inhibitory (hM4Di) or stimulatory (hM3Dq) DREADDs in the PFC-like **(e.)** or VTA-like **(f.)** component of assembloids. (For **(b,c.)**, n=8-12 per group over three separate experiments; For **(d-f.)** n=4-9 per group over 2 experiments. Data from **(c.)** is represented as mean and from **(e,f.)** is represented as mean plus individual values before and after CNO was used to activate DREADDs. Data from **(e,f.)** was analyzed with paired t-tests * p < 0.05, **p<0.01, ***p<0.001, ****p<0.0001

We modeled neural circuit-specific projections by pairing a PFC- and VTA-like spheroid together such that one expressed GCaMP6f and the other expressed the inhibitory DREADDs virus, hM4Di (**Fig. 8d**). We first recorded from assembloids where the VTA-like component of the assembloid expressed GCaMP6f and the PFC-like component expressed hM4Di and recorded both baseline 60-min after CNO activity (**Fig. 8d, left image, 8e, S8c)**. Inhibiting the PFC-like component of the assembloid significant reduced peak count to the level of a single VTA-like spheroid (**Fig. 8d,e, S8c**). Conversely, stimulating the PFC-like component increased peak count from baseline, though this was not statistically significant (**Fig. 8e, S8e**). Furthermore, baseline peak amplitude and width, along with synchrony were not changed after inhibiting the PFC-like component of the assembloid, but a significant reduction across these three parameters was observed after stimulating the PFC-like component (**Fig. 8e, S8c,e**).

We then recorded from an assembloid where the PFC-like component expressed GCaMP6f and the VTA-like component expressed either hM4Di or hM3Dq DREADDs (**Fig. 8f,g, S8d**). Inhibiting the VTA-like component of the assembloid significantly increased peak count from baseline, though this was not observed when stimulating the VTA-like component of the assembloid (**Fig. 8d,f, S8d,f**). Conversely, peak amplitude was not affected after inhibiting the VTA-like component of the assembloid but was significantly reduced after stimulating the VTA-like component, while peak width was unaffected by both stimulation or inhibition (**Fig. 8d,f, S8d,f**). Lastly, we saw that synchrony was increased after inhibiting the VTA-like component of the assembloid, though this was only a statistical trend toward significance, but significantly reduced after stimulating the VTA-like component (**Fig. 8d,f, S8d,f**). Taken together, this data shows that brain region-specific neural circuits can be modeled with these spheroids, and that calcium activity phenotypes can be modulated by manipulating different components of the assembloid.

## DISCUSSION

Three-dimensional (3D) iPSC-derived brain organoids are being used as *in vitro* models of brain development and neurological diseases (Di Lullo and Lancaster et al., 2013; Kriegstein, 2017; Pasca et al., 2018; Qian et al., 2019). Furthermore, neural circuit modeling has previously been established via the fusion of two or more different organoid types to create functional assembloids, which can improve models for neurological diseases impacting specific neural circuits such as substance use disorder (Andersen et al., 2020; Birey et al., 2017; Miura et al., 2020; Pasca et al., 2018). While organoid models have made significant inroads as 3D neural models, their complexity hinders their use as robust high-throughput drug screening (HTS) assay platforms (Andrews and Kriegstein, 2022), including batch-to-batch variation in both size and cell composition heterogeneity, limited differentiation of neuronal cell types, and lengthy differentiation and maturation times (Andrews and Kriegstein, 2022; Quadrato et al., 2016). Spheroids have emerged as physiologically relevant functional neural 3D tissue models with cell type complexity, albeit without the tissue-like anatomy, and with the robustness and reproducibility necessary for HTS. Available neural spheroid models have been generated by differentiating spheroids of neural stem cells into neurons and glial subtypes but are limited to modeling cortical brain regions (Nzou et al., 2018; Slavin et al., 2021; Woodruff et al., 2020). We expanded on this by developing a protocol to create brain region-specific neural spheroids by aggregation of different pre-differentiated neural cells, in ratios that reflected those in different regions of the human brain. The protocol developed produced spheroids homogenous in size and cell distribution, without a necrotic core, and with a final neuronal type composition similar to that initially seeded. After a 3-week maturation period, spheroids were active as shown through synaptogenesis and spontaneous intracellular calcium oscillations. We observed phenotypic profiles between single neuron spheroids that were more distinctly unique when they contained astrocytes. Given the vital role astrocytes play in synaptic connections and neurological disease (Chung et al., 2015; Lee et al., 2022; Ota et al., 2013), we generated all neuronal spheroids with a ratio of 90% neurons and 10% astrocytes.

In our study, we have shown that calcium activity can be measured using either a calcium dye (Cal6) or genetically encoded calcium indicator (GCaMP6f) as a primary readout for neuronal spheroid activity under healthy and diseased conditions. We recorded both image-based single-cell recordings with an automated confocal microscope and population, well-based calcium fluorescence measurements with a fluorescent imaging plate reader (FLIPR), a HTS instrument that measures fluorescence simultaneously across all wells on a multiwell plate. We used a multiparametric analysis to quantitate calcium activity peak features and measure differences between spheroids of different composition or with disease mutations. We applied a machine learning approach to quantify predictability of calcium activity phenotypes under healthy or diseased conditions by using a random forest classifier model and were able to demonstrate these spheroids as a reproducible endpoint to establish the effects of drugs on correcting two different disease phenotypes. While the population-based readout with FLIPR limited the detection of activity in spheroids with high GABAergic neurons, the single cell image-based analysis was able to quantitate synchronicity of the neural network, which became an important additional parameter to distinguish activity in spheroids of different composition and disease mutations, as well as corrective effects of drug treatments.

For this study, we focused on two specific brain regions, and created spheroids with neuronal subtype distributions modeling the human prefrontal cortex (PFC-like spheroids) and ventral tegmental area (VTA-like spheroids), given the role that these two brain regions play in neurological diseases including opioid use disorder (OUD), Parkinson’s Disease, and Alzheimer’s Disease (AD). The use of patient-derived cells or genetically modified cells relevant to specific neurological diseases will be critical towards the develop of personalized cellular models for precision medicine. As such, we first developed a disease model for AD by incorporating genetically engineered GABA neurons carrying the apolipoprotein e4/4 allele, which is the strongest genetic risk factor for developing AD in humans, into PFC-like spheroids given that AD leads to neurodegenerative of neocortical regions (Di Battista et al., 2016). Symptoms of AD such as altered cognitive functioning and information processing are associated with altered GABAergic signaling (Prevot and Sibillie, 2021), and correspondingly, we found baseline disruptions in peak count and width in PFC-like spheroids with APOE4/4 GABA neurons. In line with data from animal studies finding GABA-mediated disruptions in synchronous neural activity between the PFC and hippocampus, we also found that APOE4/4 GABA neurons significantly reduced synchronous neural activity compared to wildtype (Wt) control spheroids, suggesting altered GABAergic transmission (Booker et al., 2020; Fee et al., 2021; Prevot and Sibillie, 2021). Using a similar approach, we incorporated A53T alpha-synuclein dopaminergic neurons into VTA-like spheroids given that the A53T mutation is associated with early onset familial PD (Dorszewska et al., 2022). In VTA-like spheroids with A53T dopaminergic neurons, we observed increased peak count, reduced peak amplitude and width. Additionally, these spheroids showed a significant increase in the number of subpeaks, which may have resulted from increased glutamatergic and GABAergic transmission as a compensatory mechanism from loss of function in the A53T mutant neurons (Alberico et al., 2015). In line with this hypothesis, we also observed a loss of synchronous neural activity in VTA-like spheroids with A53T dopaminergic neurons, suggesting altered GABAergic transmission.

To be able to use 3D organotypic assays for disease modeling and drug discovery, it is critical to establish their context of use, that is, what stages and mechanisms underlying diseases are reported in a particular assay system, and therefore, what targeted pharmacologically agents will perturb the activity of the system (Eckert et al. 2020, Jung et al. 2021). Here, we used a panel of drugs that are being used to treat symptoms of AD or PD, with different mechanisms of action, to assess what targeted pharmacological perturbations reversed the disease phenotypes in our disease spheroid models. For AD, we tested five compounds, and saw that three reversed the AD disease phenotypes, including Memantine and Donepezil, two clinically approved compounds used to treat AD, and EUK-134, a preclinical compound. While these compounds treat the symptoms of AD rather than the underlying disease, a recent meta-analysis showed that Memantine has shown credible efficacy when prescribed alone or in combination with cholinesterase inhibitors such as Donepezil (Kishi et al., 2017). We also found that EUK-134, which has been associated with reduced beta-amyloid plaques in mouse models of AD (Bahramikia and Yazdanparast, 2013; Jekabsone et al., 2006), was able to reverse deficits caused by APOE 4/4 GABA neurons in our PFC-like AD neuronal spheroid model. For the PD model, we tested eight compounds, and only Ropinirole, a dopamine agonist, rescued deficits in dopaminergic transmission, suggesting that pharmacological response may be patient-but also mutation-specific. Further studies are under way in both disease models to assess the presence of clinical hallmarks of AD and PD in each respective model to further establish its context of use and mechanistically explain the different pharmacological responses obtained by each drug.

To model OUD, we tested both PFC- and VTA-like spheroids given the role these two brain regions play in drug reward, withdrawal, and relapse (Koob, 2021). PFC- and VTA-like spheroids were chronically treated with DAMGO, a μ-opioid receptor (MOR) agonist, for ten days prior to calcium activity recordings, with and without a 3-day washout period mimicking withdrawal. In line with clinical imaging studies showing alterations in magnetic resonance imaging (MRI) signal in the PFC and VTA of people with opioid use disorder (Goldstein and Volkow, 2011; Volkow et al., 2019), we found changes in baseline activity in both spheroid types, where chronic DAMGO treatment in PFC-like spheroids reduced peak count while in VTA-like spheroids chronic DAMGO and DAMGO withdrawal increased peak count. Interestingly, treatment with naloxone, a MOR antagonist used to reverse opioid overdoses in humans, reversed DAMGO-induced reductions in peak count in PFC-like spheroids while having no impact on VTA-like spheroids, suggesting that the mechanism may be through glutamatergic and GABAergic transmission more than dopaminergic in these spheroids.

In conclusion, we have developed a highly reproducible functional neural spheroid assay platform where cell type composition can be adjusted to mimic a specific brain region of interest and that can be used for HTS and neural circuitry modeling. We also developed disease models for AD, PD, and OUD, and machine learning classifier models were used to quantify predictability of disease phenotypes, showing that the AD and PD model showed highly predictive disease-related phenotypes. In addition, we show that our methodology can be used to fuse neuronal spheroids to create assembloids and establish neural circuitry. Future studies aim to further establish the context of use of these neural disease models for drug discovery and development by doing a more in depth phenotypic and genomic analysis of disease biomarkers, pharmacological profiling with additional compounds of relevant targets and mechanisms and develop high-throughput methods for making neural circuit-specific assembloids to study circuit-level effects of compounds on neurological diseases. Overall, the current study lays the groundwork for potentially improving drug discovery for neurological diseases by creating physiological and pathological relevant tissue models that are robust and have the throughput for drug discovery and development.

## AUTHOR CONTRIBUTIONS

C.E.S., S.K., M.E.B., and E.L. performed experiments. C.E.S., Y.C., and K.W. performed analysis. C.E.S., E.L., and M.F. designed experiments and planned study designs. All authors participated in writing and/or editing the manuscript.

## ACKNOWLEDGEMENTS

This research was supported by the Helping End Addiction Longterm (HEAL) NIH program at the National Center for Advancing Translational Sciences.

