## Supplemental Table 1 for "Functional brain region-specific neural spheroids for modeling neurological diseases and therapeutics screening"

| VTA-like | Plate 1 | Plate 2 | Plate 3 | Plate 4 | Plate 5 | Plate 6 |
|---|---|---|---|---|---|---|
| Peak Amplitude (RFU) | 14 | 13 | 12 | 21 | 20 | 13 |
| Peak Amplitude SD | 131 | 93 | 78 | 43 | 108 | 45 |
| Peak Count | 12 | 12 | 14 | 11 | 9 | 15 |
| Peak Rate (s) | 12 | 17 | 16 | 10 | 8 | 13 |
| Peak Rate SD | 143 | 150 | 112 | 69 | 85 | 65 |
| Peak Spacing (s) | 10 | 11 | 13 | 9 | 5 | 13 |
| Peak Spacing SD | 70 | 59 | 56 | 40 | 42 | 39 |
| Peak Width 50% | 12 | 14 | 15 | 12 | 7 | 15 |
| Peak Width 90% | 10 | 11 | 13 | 10 | 6 | 13 |
| Rise Slope (RFU) | 29 | 23 | 28 | 30 | 29 | 25 |
| Rise Slope SD | 54 | 51 | 51 | 51 | 38 | 42 |
| Peak Rise Time (s) | 28 | 19 | 28 | 18 | 20 | 16 |
| Peak Rise Time SD | 118 | 84 | 109 | 79 | 46 | 59 |
| Decay Slope (RFU) | 17 | 18 | 21 | 23 | 21 | 24 |
| Decay Slope SD | 46 | 50 | 46 | 44 | 53 | 42 |
| Peak Decay Time (s) | 12 | 16 | 16 | 11 | 4 | 26 |
| Peak Decay Time SD | 47 | 99 | 49 | 42 | 35 | 54 |

| PFC-like | Plate 1 | Plate 2 | Plate 3 | Plate 4 | Plate 5 | Plate 6 |
|---|---|---|---|---|---|---|
| Peak Amplitude (RFU) | 16 | 18 | 17 | 35 | 25 | 10 |
| Peak Amplitude SD | 59 | 51 | 30 | 35 | 30 | 17 |
| Peak Count | 11 | 24 | 10 | 20 | 22 | 15 |
| Peak Rate (s) | 12 | 21 | 10 | 19 | 23 | 15 |
| Peak Rate SD | 40 | 31 | 18 | 43 | 42 | 19 |
| Peak Spacing (s) | 12 | 35 | 11 | 29 | 21 | 16 |
| Peak Spacing SD | 34 | 94 | 25 | 51 | 23 | 40 |
| Peak Width 50% | 17 | 37 | 10 | 38 | 16 | 15 |
| Peak Width 90% | 12 | 28 | 8 | 31 | 9 | 14 |
| Rise Slope (RFU) | 24 | 26 | 15 | 55 | 26 | 12 |
| Rise Slope SD | 21 | 27 | 17 | 37 | 28 | 14 |
| Peak Rise Time (s) | 43 | 39 | 7 | 70 | 39 | 11 |
| Peak Rise Time SD | 154 | 174 | 9 | 115 | 166 | 24 |
| Decay Slope (RFU) | 15 | 20 | 14 | 39 | 21 | 12 |
| Decay Slope SD | 47 | 42 | 14 | 37 | 41 | 12 |
| Peak Decay Time (s) | 11 | 32 | 8 | 18 | 8 | 13 |
| Peak Decay Time SD | 97 | 97 | 22 | 75 | 78 | 31 |

| *Plate 7* | *Plate 8* | *Plate 9* | *Plate 10* | *mean* |
|---|---|---|---|---|
| 13 | 24 | 11 | 19 | 16 |
| 69 | 46 | 23 | 193 | 83 |
| 15 | 17 | 10 | 15 | 13 |
| 19 | 29 | 7 | 10 | 14 |
| 131 | 120 | 53 | 102 | 103 |
| 11 | 17 | 6 | 10 | 11 |
| 91 | 84 | 45 | 74 | 60 |
| 13 | 22 | 8 | 10 | 13 |
| 7 | 25 | 6 | 7 | 11 |
| 33 | 24 | 21 | 21 | 26 |
| 76 | 70 | 37 | 60 | 53 |
| 44 | 31 | 18 | 17 | 24 |
| 101 | 133 | 62 | 82 | 87 |
| 23 | 40 | 15 | 21 | 22 |
| 44 | 62 | 41 | 53 | 48 |
| 29 | 58 | 11 | 18 | 20 |
| 34 | 128 | 32 | 56 | 58 |

| *Plate 7* | *Plate 8* | *Plate 9* | *Plate 10* | *mean* |
|---|---|---|---|---|
| 19 | 17 | 21 | 11 | 19 |
| 41 | 47 | 38 | 41 | 39 |
| 19 | 29 | 18 | 6 | 18 |
| 19 | 29 | 18 | 7 | 17 |
| 25 | 49 | 19 | 81 | 37 |
| 19 | 41 | 20 | 8 | 21 |
| 45 | 69 | 42 | 164 | 59 |
| 14 | 40 | 18 | 7 | 21 |
| 12 | 27 | 15 | 5 | 16 |
| 21 | 20 | 20 | 26 | 24 |
| 35 | 33 | 23 | 19 | 25 |
| 13 | 19 | 8 | 31 | 28 |
| 15 | 119 | 15 | 100 | 89 |
| 12 | 16 | 13 | 17 | 18 |
| 25 | 61 | 27 | 42 | 35 |
| 7 | 20 | 13 | 9 | 14 |
| 28 | 85 | 29 | 61 | 60 |
