## Supplemental Information for "Functional brain region-specific neural spheroids for modeling neurological diseases and therapeutics screening"

**Supplemental Results**

**Designer neural spheroids exhibit high well-to-well reproducibility**

Baseline recordings were collected with the fluorescent imaging plate reader (FLIPR) for all plates tested prior to the addition of compounds with a 384-well pin tool. For all recordings, spheroids incubated in Cal6 dye for 2-hrs before the baseline recording. For FLIPR recordings, a multiparametric approach was used for analysis, and for each plate tested, 17 peak parameters were measured using Molecular Devices ScreenWorks PeakPro 2.0 peak detection analysis. Given that these spheroids were being tested for the high-throughput screening (HTS) capabilities, coefficients of variance (CV) values were calculated from data obtained from baseline FLIPR recordings to determine which of the 17 peak parameters were reproducible and, therefore, used across all plates and recordings. CV values were calculated by dividing the standard deviation over the mean for each peak parameter analyzed within each spheroid type, and values were reported as percentages (%CV). Out of the 17 peak parameters measured, 10 consistently showed %CV values ≤30%, indicating high well-to-well reproducibility. Table S1 shows the %CV values for each plate along with the average across all plates tested, with the 10 peak parameters ≤30% including peak count, peak rate, peak spacing, peak width 50% and 90%, peak amplitude, peak rise time, peak decay time, rise slope, and decay slope (**Table S1**).

**Designer spheroids express neuronal cell-type markers for dopaminergic, glutamatergic, and GABAergic neurons in quantities reflecting their cell-type composition**

Spheroids were stained for tyrosine hydroxylase (TH), vesicular glutamate transporter 1 (vGluT1), and parvalbumin (PV) to label dopaminergic, glutamatergic, and GABAergic neurons, respectively. Image stacks were collected on a confocal microscope through the entirety of a “VTA-like” and “PFC-like” spheroid. Videos going through each z-stack were made to display expression of each cell-type marker, and a video showing the “VTA-like” spheroid is shown in **Video S1** while the “PFC-like” spheroid is shown in **Video S2**. In both spheroid types, Hoechst nuclear expression is distributed evenly throughout the entire spheroid, suggesting these spheroids lack a necrotic core (**Video S1, S2**). Furthermore, “VTA-like” spheroids, which contain 65% dopaminergic neurons, showed that the majority of neurons were labeled with TH (**Video S1**). While “PFC-like” spheroids were not generated with dopaminergic neurons, a few cells within this spheroid type expressed TH, which may be reflective of neuronal cells that are not 100% pure prior to spheroid generation (**Video S2**). Both spheroid types expressed vGluT, though this expression was higher in “PFC-like” spheroids, and both types showed similar expression of PV given that each spheroid type contained 30% GABA neurons (**Video S1, S2**). vGluT1 is a presynaptic marker and immunostaining for this labeled whole neurons, we followed up with a FISH assay to examine mRNA expression of vGluT in each spheroid type. Using FISH, we observed puncta expression of vGluT, in line with a synaptic marker (**Fig. S1**). Additionally, vGluT expression was significantly lower in “VTA-like” spheroids, which are generated with 5% glutamatergic neurons, compared to “PFC-like” spheroids, which are generated with 70% glutamatergic neurons (**Fig. S1**).

**Astrocytes are not necessary for neuronal activity but alter phenotypic profiles in spheroids containing dopaminergic and glutamatergic neurons**

Prior to developing brain region-specific spheroids, single neuron spheroids (SNSs) were compared with and without astrocytes to understand the contribution of astrocytes to neuronal activity. Here, SNSs containing dopaminergic, glutamatergic, or GABAergic neurons were either generated as spheroids containing 100% of their respective neuronal subtype or containing 90% neuronal subtype and 10% astrocytes. Calcium activity was recorded with an automated confocal and peak parameters including peak count, amplitude, and width were measured at the individual cell level using the automated ROI identifier, LC_Pro, along with synchrony across all identified ROIs (**Fig. S2**). Reduced peak count was observed in in SNSs with dopaminergic neurons and astrocytes but increased in glutamatergic SNSs with astrocytes (**Fig. S2b,** left panel; unpaired t-tests, dopaminergic: t_(8)_=5.02, p=0.001; glutamatergic: t_(7)_=5.29, p=0.001; GABAergic: t_(8)_=0.046, p=0.96). Conversely, increased peak amplitude and width were observed in dopaminergic SNSs with astrocytes compared to those without astrocytes while the opposite was observed in SNSs with glutamatergic neurons (**Fig. S2b,** middle panels; unpaired t-tests for peak amplitude and width, respectively; dopaminergic: t_(7)_=8.56, p<0.0001, t_(8)_=7.71, p<0.0001; glutamatergic: t_(6)_=9.45, p<0.0001, t_(7)_=7.28, p=0.0002; GABAergic: t_(8)_=1.004, p=0.35, t_(8)_=1.41, p=0.195). In SNSs with dopaminergic and GABAergic neurons, astrocytes did not impact neuronal synchrony, however there was a significant reduction in synchrony among SNSs containing glutamatergic neurons and astrocytes compared to those without astrocytes (**Fig. S2b,** right panel; dopaminergic: t_(8)_=2.13, p=0.066; glutamatergic: t_(7)_=2.76, p=0.028; GABAergic: t_(8)_=0.959, p=0.37).

**Functional responses to quality control (QC) compounds are different by spheroid type**

Functional responses within each spheroid type were validated via treatment with QC compounds targeting receptors for each neuronal subtype. Immediately after the baseline recording, spheroids were treated with QC compounds and responses were measured 1-,30-, and 60-min later. Linear mixed model (LMM) ANOVA was used to examine repeated measures, with treatment as the between-subjects factor and time as the within-subjects factor. Within each spheroid type, two-way LMM ANOVAs were performed for each peak parameter measured, and significant main effects were observed across all peak parameters for both spheroid types (“VTA-like” spheroids: peak amplitude: F_(24,222)_=39.55, p<0.0001, peak count: F_(24,222)_=67.25, p<0.0001, peak spacing: F_(24,222)_=39.28, p<0.0001, peak rate: F_(24,222)_=15.99, p<0.0001, peak width 50%: F_(24,222)_=40.1, p<0.0001, peak width 90%: F_(24,222)_=19.82, p<0.0001, rise slope: F_(24,222)_=37.08, p<0.0001, peak rise time: F_(24,222)_=28.26, p<0.0001, decay slope: F_(24,222)_=39.23, p<0.0001, peak decay time: F_(24,222)_=18.06, p<0.0001; “PFC-like” spheroids: peak amplitude: F_(21,253)_=51.85, p<0.0001, peak count: F_(21,253)_=30.4, p<0.0001, peak spacing: F_(21,253)_=27.03, p<0.0001, peak rate: 44.5, p<0.0001, peak width 50%: F_(21,253)_=34.08, p<0.0001, peak width 90%: F_(21,253)_=45.23, p<0.0001, rise slope: F_(21,253)_=37.23, p<0.0001, peak rise time: F_(21,253)_=41.69, p<0.0001, decay slope: F_(21,253)_=54.72, p<0.0001, peak decay time: F_(21,253)_=7.95, p<0.0001). Significant main effects were followed up with Tukey’s posthoc and all statistical data is reported in **Table S2**, with significant values below 0.05 in bold.

In both spheroid types, GABA_A_R agonism with Muscimol treatment led to a total inhibition that occurred 1-,30-, and 60-min after treatment, despite the fact that no differences were observed from DMSO-treated controls during the baseline recording (**Fig. S3, Table S2**). Treatment with GABA_A_R antagonist, Bicuculline induced differential functional changes between the two spheroid types, including changes such as an increase in peak count and decrease in peak amplitude in “VTA-like” spheroids but the opposite in “PFC-like” spheroids, both of which began during 1-min after treatment and persisted 60-min later (**Fig. S3, Table S2**). In both spheroid types, blocking NMDARs with Memantine led to a total inhibition of activity 1-min after treatment, though 30- and 60-min after treatment both spheroid types displayed enhanced stimulatory activity, as evidenced by changes including significantly increased peak count and rate but decreased peak spacing (**Fig. S3, Table S2**). Interestingly, blocking AMPARs with CNQX led to a total inhibition that persisted across all three recordings in “PFC-like” spheroids, but this inhibition was only observed in “VTA-like” spheroids 1-min after treatment, and while some peak parameters such as peak rise time remained significantly higher 30- and 60-min after treatment, peak count and spacing were the same as DMSO-treated controls (**Fig. S3, Table S2**). In both spheroid types, blocking dopamine 1 receptors (D1Rs) with SCH23390 inhibited activity 1-,30-, and 60-min after treatment (**Fig. S3, Table S2**). In “VTA-like” spheroids, blocking D2Rs with sulpiride primarily altered peak width and peak decay time, which did not begin until 30-min after treatment and was also observed 60-min after treatment (**Fig. S3, Table S2**). However, in “PFC-like” spheroids, D2R antagonism led to enhanced excitability, as indicated by significant increases in peak count and rate but reduced peak spacing, though these were not significantly different from DMSO-treated controls until 60-min after treatment (**Fig. S3, Table S2**).

**Neurological disease models for Alzheimer’s and Parkinson’s Disease as well as Opioid Use Disorder do not impact spheroid viability**

To understand whether functional differences observed across each disease model could be attributed to differences in spheroid viability, the 3D Cell Titer Glo assay was used to assess spheroid viability in a subset of spheroids that were not treated with compounds (**Fig. S4**). For the Alzheimer’s Disease (AD) model with apolipoprotein e4/4 (APOE4/4) GABA neurons, viability was measured in “PFC-like” and SNSs with wildtype (Wt) or APOE4/4 GABA neurons. Unpaired t-tests showed no significant differences in viability for the GABAergic SNSs as well as “PFC-like” spheroids (**Fig. S4a,** SNSs: t_(16)_=1.82, p=0.087; “PFC-like”: t_(17)_=0.587, p=0.56). For the Parkinson’s Disease (PD) model with mutant A53T alpha-synuclein dopaminergic neurons, viability was measured in “VTA-like” and SNSs with Wt or A53T dopaminergic neurons. Unpaired t-tests showed no significant differences in viability for the dopaminergic SNSs as well as “VTA-like” spheroids (**Fig. S4b,** SNSs: t_(14)_=0.365, p=0.72; “VTA-like”: t_(13)_=0.39, p=0.71). One way ANOVA was used to measure spheroid viability in both “PFC-like” and “VTA-like” spheroids receiving no DAMGO pre-treatment, chronic DAMGO pre-treatment, or chronic DAMGO pre-treatment with a 3-day washout period to mimic withdrawal (WD), and no significant differences were observed in either spheroid type (**Fig. S4c,** “PFC-like”: F_(2,26)_=3.07, p=0.064; “VTA-like”: F_(2,28)_=3.03, p=0.064).

**Effects of compounds used to treat spheroids modeling Alzheimer’s Disease (AD) on Wt “PFC-like” spheroids**

For the AD model, we also assessed the effect of compounds tested on APOE4/4 “PFC-like” spheroids in Wt “PFC-like” spheroids to understand if the effects they induced were specific to mutant spheroids. At baseline, the only group with significantly different peak count and spacing from Wt DMSO-treated controls was DMSO-treated APOE4/4 “PFC-like” spheroids (**Fig. S5,** One way ANOVA: Peak count: F_(8,94)_=13.65, p<0.0001, Dunnett’s posthoc: Wt DMSO vs APOE4/4 DMSO, t_(94)_=6.33, p<0.0001; Peak spacing: F_(8,94)_=20.17, p<0.0001, Dunnett’s posthoc: Wt DMSO vs APOE4/4 DMSO, t_(94)_=8.52, p<0.0001). 90-min after treatment, significant differences in peak count from DMSO-treated controls in Wt “PFC-like” spheroids occurred after treatment with Donepezil, Memantine, EUK-134, and Hu-210 (**Fig. S5a,** F_(8,88)_=28.82,p<0.0001; Dunnett’s posthoc: Wt DMSO vs 1 uM Donepezil: t_(88)_=3.62, p = 0.004; Wt DMSO vs 10 uM Donepezil: t_(88)_=7.39, p<0.0001; Wt DMSO vs 10 uM Memantine: t_(88)_=4.49, p=0.0002; Wt DMSO vs 10 uM EUK-134: t_(88)_=3.25,p=0.011; Wt DMSO vs Hu-210: t_(88)_=4.55, p=0.0001). Opposite differences in peak spacing were observed in all groups except Wt “PFC-like” spheroids treated with Rivastigmine (**Fig. S5b,** F_(8,79)_=3.32, p=0.003; Dunnett’s posthoc: Wt DMSO vs APOE4/4 DMSO: t_(79)_=9.06, p<0.0001; Wt DMSO vs 1 uM Donepezil: t_(79)_=3.51, p = 0.005; Wt DMSO vs 10 uM Donepezil: t_(79)_=4.98, p<0.0001; Wt DMSO vs 10 uM Memantine: t_(79)_=4.71, p<0.0001; Wt DMSO vs 1 uM EUK-134: t_(79)_=3.31, p=0.0098; Wt DMSO vs 10 uM EUK-134: t_(79)_=3.17, p=0.015; Wt DMSO vs Hu-210: t_(79)_=9.47, p<0.0001). Together, this data shows that these compounds, some of which were able to reverse deficits in APOE4/4 “PFC-like” spheroids have similar effects on Wt “PFC-like” spheroids.

**Effects of compounds used to treat spheroids modeling Parkinson’s Disease (PD) on Wt and A53T “VTA-like” spheroids**

In A53T mutant “VTA-like” spheroids modeling PD, we found that Ropinirole, dopamine agonist, reversed deficits in increased peak count and subpeaks to the level of Wt DMSO-treated “VTA-like” control spheroids (**Fig. 6**). However, we also tested the effects of other clinically approved drugs used to treat PD including: L-Dopa, Amantadine, Rasagiline, Entacapone, Tolcapone, Trihexyphenidyl, and Benztropine. In Wt “VTA-like” spheroids, one way ANOVA showed a significant main effect during the baseline recording, which revealed that the only group with significantly different peak count compared to Wt DMSO-treated control spheroids was in A53T DMSO-treated spheroids (**Fig. S6a,** F_(8,93)_=8.28, p<0.0001; Dunnett’s posthoc, Wt DMSO vs A53T DMSO: t_(93)_=6.11, p<0.0001). Wt “VTA-like” spheroids were then treated with compounds, and 90-min later, a significant main effect from the one-way ANOVA showed significant differences in peak count in A53T DMSO-treated compared to Wt DMSO-treated “VTA-like” spheroids, along with Benztropine and Trihexyphenidyl reducing peak count since they had an inhibitory effect on spheroids (**Fig. S6a,** F_(8,90)_=60.38, p<0.0001; Dunnett’s posthoc, Wt DMSO vs A53T DMSO: t_(90)_=5.85, p<0.0001; Wt DMSO vs Wt Benztropine: t_(90)_=11.59, p<0.0001; Wt DMSO vs Wt Trihexyphenidyl: t_(90)_=11.76, p<0.0001). Within A53T “VTA-like” spheroids, all groups showed significantly enhanced peak count compared to Wt DMSO-treated controls during both the baseline recording as well as 90-min after treatment (**Fig. S6b,** Baseline: F_(8,108)_=4.51, p<0.0001; Dunnett’s posthoc: t_(108)_>3.62, p<0.05; 90-min: F_(8,108)_=51.3, p<0.0001, t_(108)_> 2.75, p<0.05 across all groups).

**Effect of naloxone on “VTA-like” and “PFC-like” spheroids modeling Opioid Use Disorder**

In Fig. 7 we showed that naloxone was able to reverse deficits in peak count and peak spacing in “PFC-like” spheroids chronically treated with DAMGO, a mu opioid receptor (MOR) agonist. We also measured the effect of naloxone in spheroids modeling DAMGO withdrawal as they were chronically pre-treated with DAMGO but received a 3-day washout period prior to recording calcium activity. During the baseline recording, there was no effect of pre-treatment on peak count or peak spacing as indicated by one-way ANOVA (**Fig. S7a,b,** left panel, peak count: F_(3,51)_=2.11, p=0.11; peak spacing: F_(3,51)_=1.86, p=0.15). After the baseline recording, “PFC-like” spheroids were treated with either DAMGO or DMSO, and here we found that acute DAMGO treatment significantly reduced peak count and increased peak spacing compared to DMSO-treated control spheroids, but that DAMGO WD spheroids showed no differences (**Fig. S7a,b** middle panel, peak count: F_(3,50)_=2.54, p=0.067, Dunnett’s posthoc: Control DMSO vs Control DAMGO: t_(50)_=2.58, p=0.034; peak spacing: F_(3,51)_=3.67, p=0.018, Dunnett’s posthoc: Control DMSO vs Control DAMGO: t_(51)_=3.15, p=0.008). Following the 30-min recording, spheroids were treated with naloxone and no differences were observed in either peak count or peak spacing (**Fig. S7a,b** right panel, peak count: F_(3,51)_=2.19, p=0.1; peak spacing: F_(3,51)_=0.328, p=0.81). Together, these data show that “PFC-like” spheroids modeling DAMGO WD do not display the baseline deficits or response to treatment as “PFC-like” spheroids pre-treated with chronic DAMGO, suggesting chronic DAMGO pre-treatment may be a better OUD model in “PFC-like” spheroids.

In VTA-like spheroids, no significant differences were observed in either pre-treatment group during the baseline recording when examining peak count and spacing (**Fig. S7c,d,** left panel, peak count: F_(5,97)_=1.2, p=0.31; peak spacing: F_(5,97)_=1.12, p=0.35). 30-min after treatment with either DAMGO or DMSO, a significant main effect of pre-treatment indicated that spheroids pre-treated with DAMGO displayed significantly lower peak count and spacing compared to “VTA-like” spheroids in the control pre-treatment group (**Fig. S7c,d,** middle panel, peak count: F_(2,97)_=3.25, p=0.04; peak spacing: F_(2,97)_=3.49, p=0.035). However, no effects were observed 30-min after treatment with naloxone or DMSO on peak count (**Fig. S7c,** right panel, peak count: F_(2,97)_=0.39, p=0.68). Analysis of peak spacing showed a significant main effect of treatment, but multiple comparisons tests did not show any significant effects between treatment groups (**Fig. S7d,** right panel, main effect of treatment: F_(1,97)_=8.7, p=0.004). In summary, the OUD model did not lead to significant alterations in baseline activity among “VTA-like” spheroids, suggesting that “PFC-like” spheroids may be a more suitable model for OUD.

**Supplemental Figures**

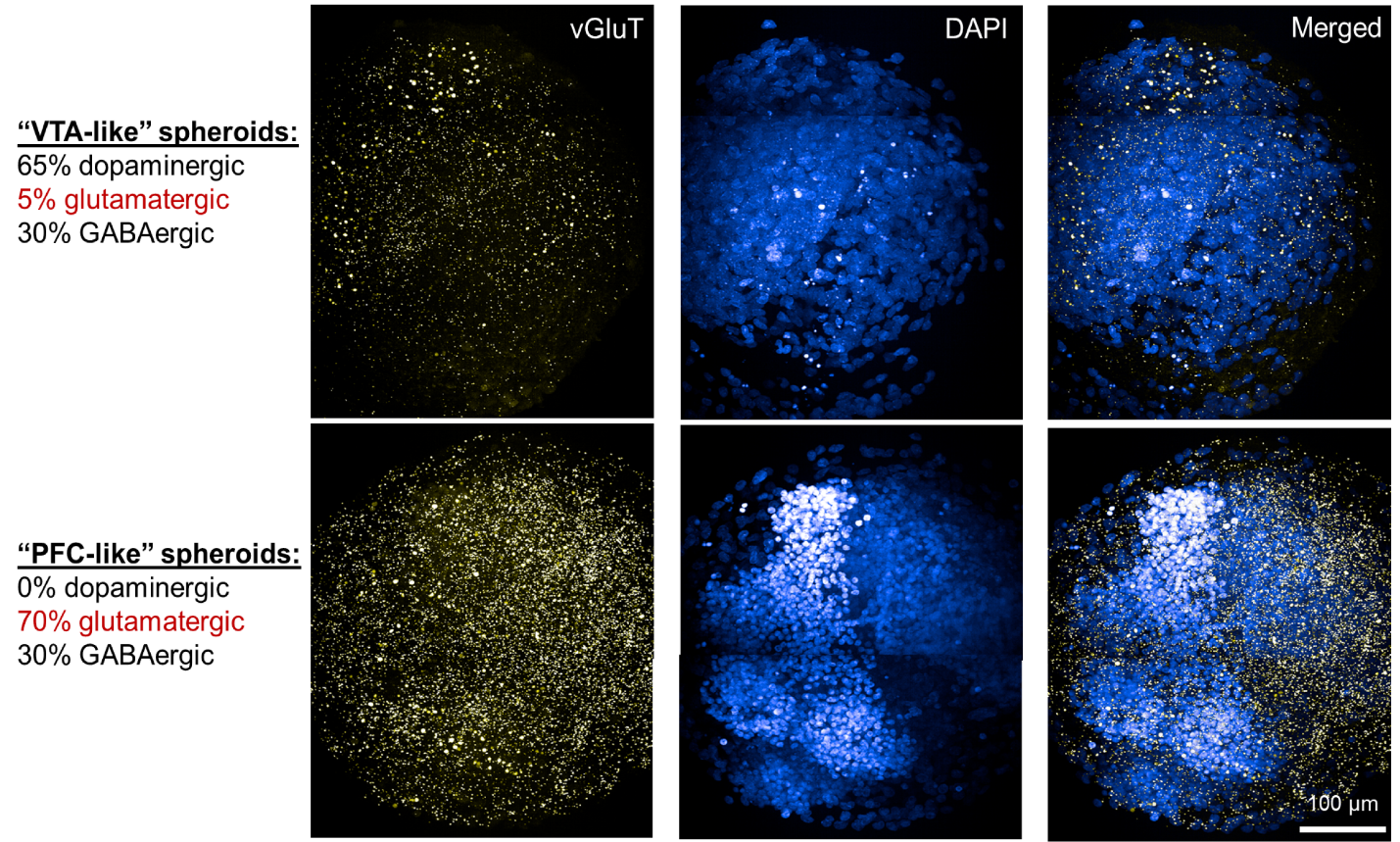

**Figure S1. vGluT mRNA expression as an indicator of glutamatergic neuron composition in “VTA-like” and “PFC-like” spheroids.** Maximum projections from a representative “VTA-like” (top panel) and “PFC-like” spheroid showing vGluT mRNA to demonstrate glutamatergic neuronal subtype composition within each spheroid. From left to right, images shown include vGluT (pseudo-colored gold), DAPI nuclear stain (pseudo-colored blue), and a merged image showing vGluT and DAPI. 350 um z-stacks were collected with a 63X oil objective, and maximum projection images across all z-planes are displayed.

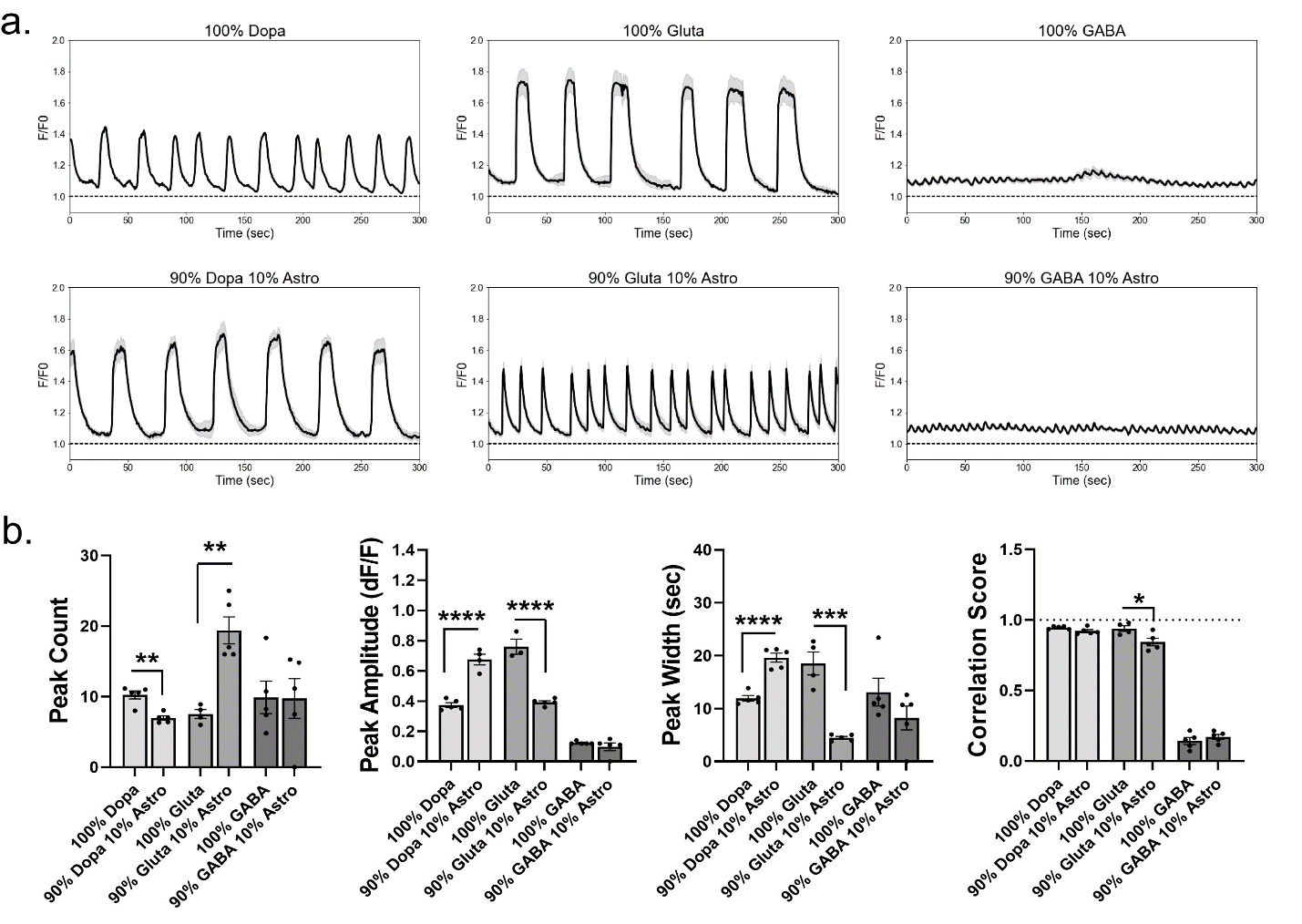

**Figure S2. Astrocytes are not necessary for neuronal activity, but their presence alters phenotypic profiles of single neuron spheroids (SNSs). (a.)** Data collected from Phenix Plus automated confocal microscope recordings obtained from spheroids in a Cal6 dye; Representative time series plots showing calcium activity phenotypes in SNSs without astrocytes (100% dopaminergic, glutamatergic, or GABAergic neurons) or with 10% astrocytes and 90% neuron **(b.)** Quantified peak count, amplitude, and width data along with correlation score representing the average R^2^ value from all identified ROIs in each spheroid. In SNS dopaminergic spheroids, astrocytes reduced peak count and increased peak amplitude/width. In SNS glutamatergic spheroids, astrocytes increased peak count and reduced peak amplitude/width along with a slight reduction in synchrony. Astrocytes did not alter activity in SNSs with GABAergic neurons. (Data was run in technical replicates of n=4-6 per plate across one experiment for SNSs; For b., unpaired t-tests were used to compare SNSs with the same neuronal subtype (with vs without astrocytes), data is represented as mean ± SEM, * p<0.05, ** p<0.01, *** p < 0.001, ****p<0.0001.

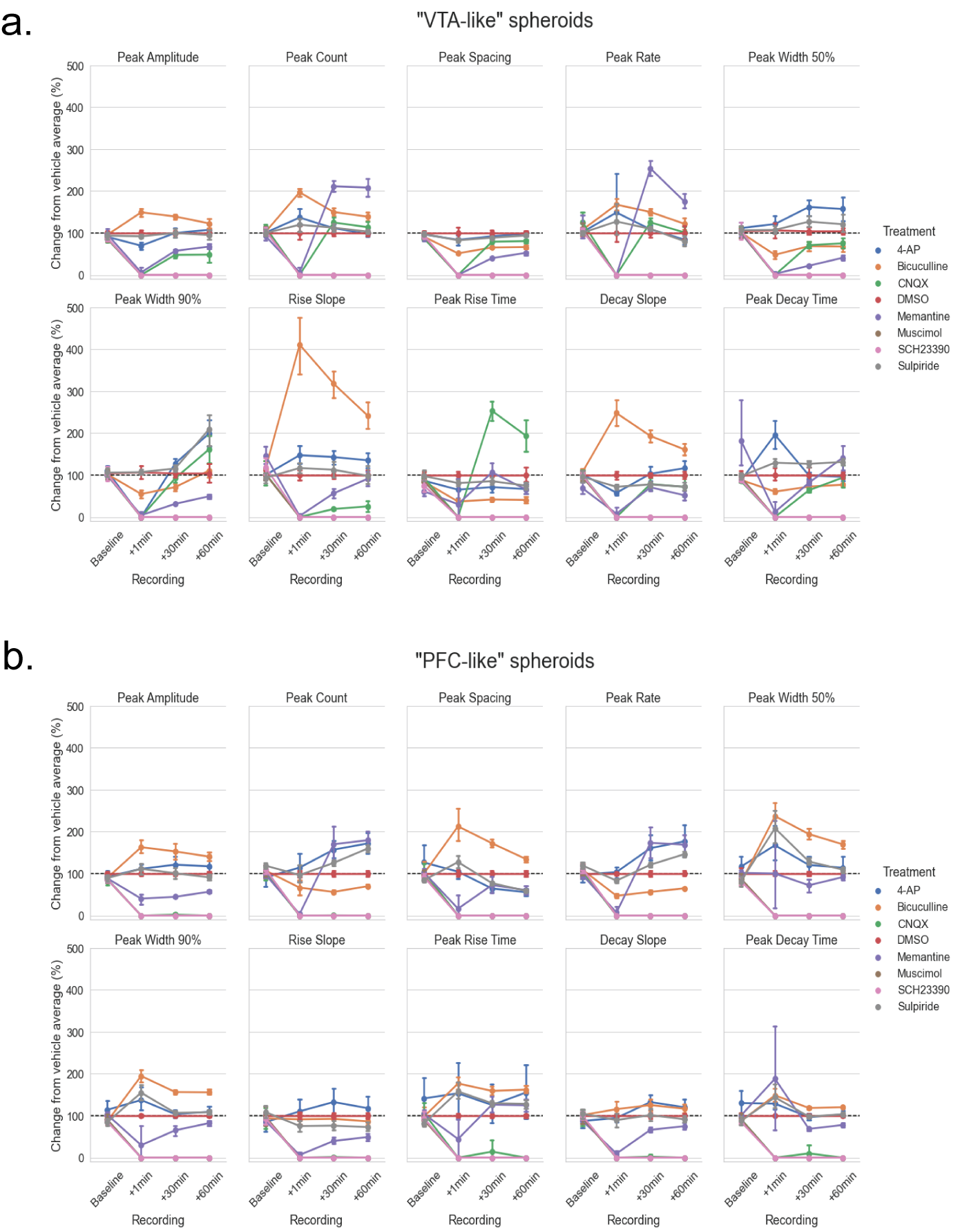

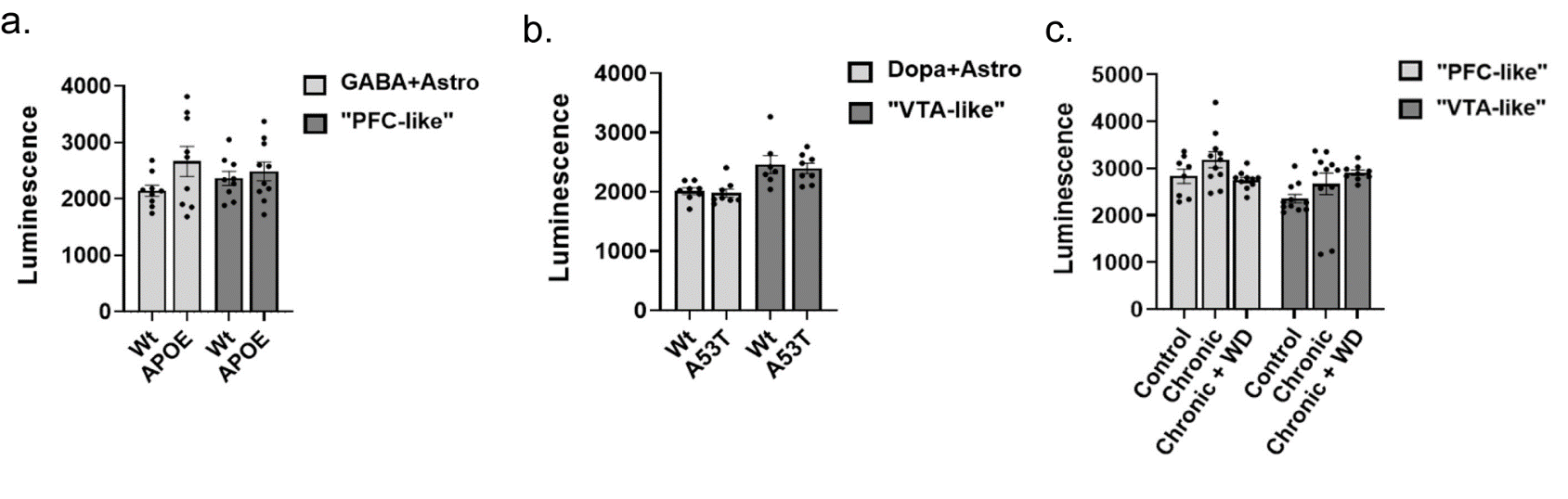
**Figure S3. Functional responses to control compounds in brain region-specific neural spheroids throughout the 60-min recording period. (a.)** Data from VTA-like spheroids **(b.)** Data from PFC-like spheroids **(a,b.)** Calcium activity, relative to DMSO-treated controls, during four recordings including baseline plus 1-, 30-, and 60-min after treatment with control compounds. Compounds used to target GABA_A_ receptors included agonist, muscimol, and antagonist, bicuculline. Ionotropic glutamate receptors were targeted with CNQX, AMPAR antagonist, and Memantine, NMDAR antagonist while dopamine receptors were targeted with SCH23390, D1R antagonist, and sulpiride, D2R antagonist. Data is shown for all 10 peak parameters analyzed and is represented as mean ± 95% confidence interval. Linear mixed model ANOVA was used to examine repeated measures fitted to a mixed model, with treatment as the between-subjects factor and recording as the within-subjects factor. Analysis results are represented in Table S2.

**Figure S4. Comparisons of spheroid viability and select peak parameters from FLIPR data between wildtype and disease models. (a-c.)** Luminescence from spheroids from the 3D Cell Titer Glo (CTG) assay to measure spheroid viability. **(a.)** Comparisons between Wt and APOE4/4 GABA neurons in single neuron spheroids (SNSs) with GABAergic neurons and “PFC-like” spheroids **(b.)** Comparisons between Wt and A53T dopaminergic neurons in SNSs with dopaminergic neurons and “VTA-like” spheroids **(c.)** Comparisons between DAMGO pre-treatment groups, including control, chronic DAMGO, and DAMGO WD in both “PFC-like” and “VTA-like” spheroids (3D CTG assay: n=8-10 technical replicates, n=1 biological replicate for each disease line collected over three independent experiments. Results are analyzed with unpaired t-tests (a,b) and One way ANOVA (c.) with significance set at p<0.05. Data from are represented
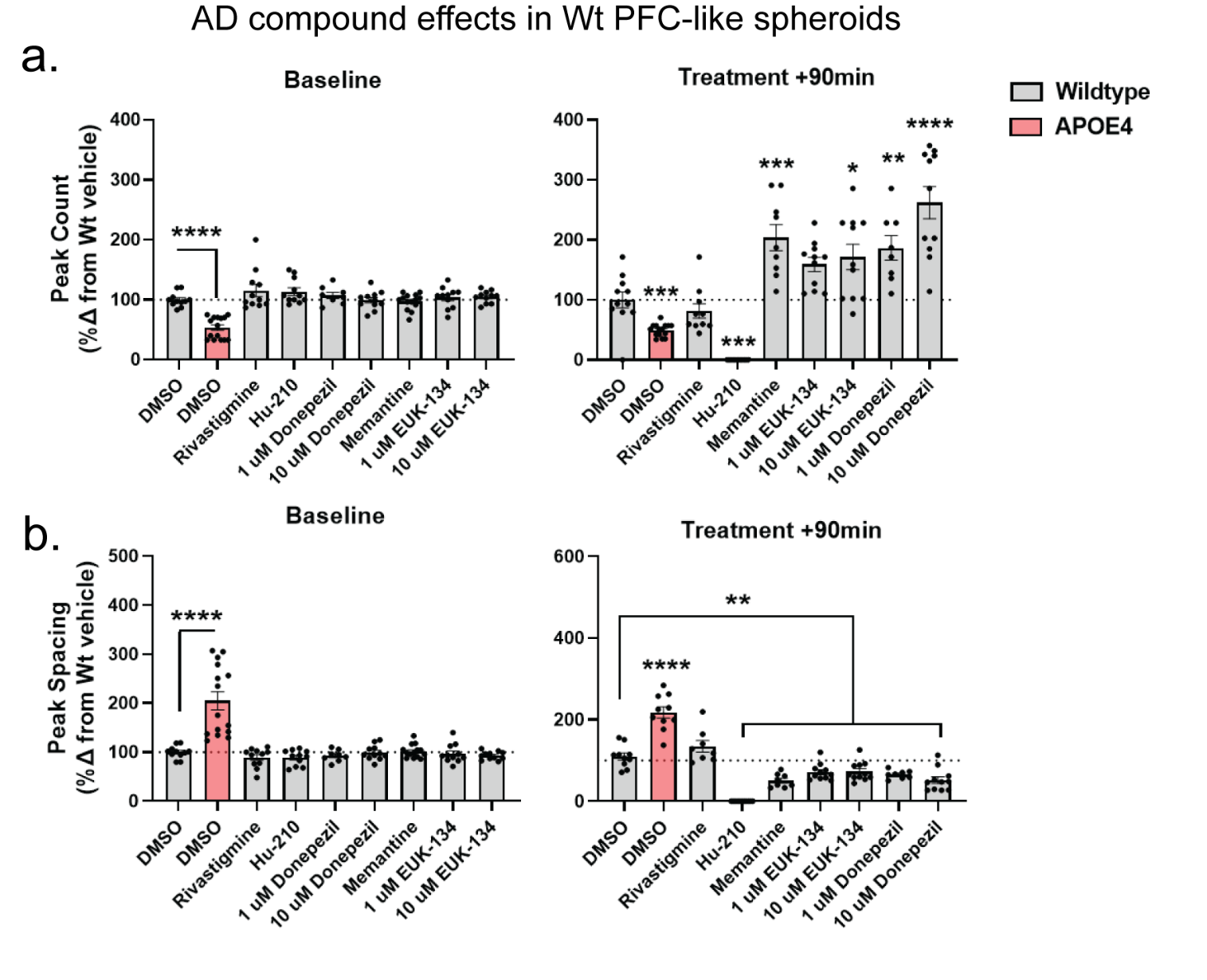
as mean ± SEM; *p<0.05, **p<0.01, ***p<0.001, ****p<0.0001.

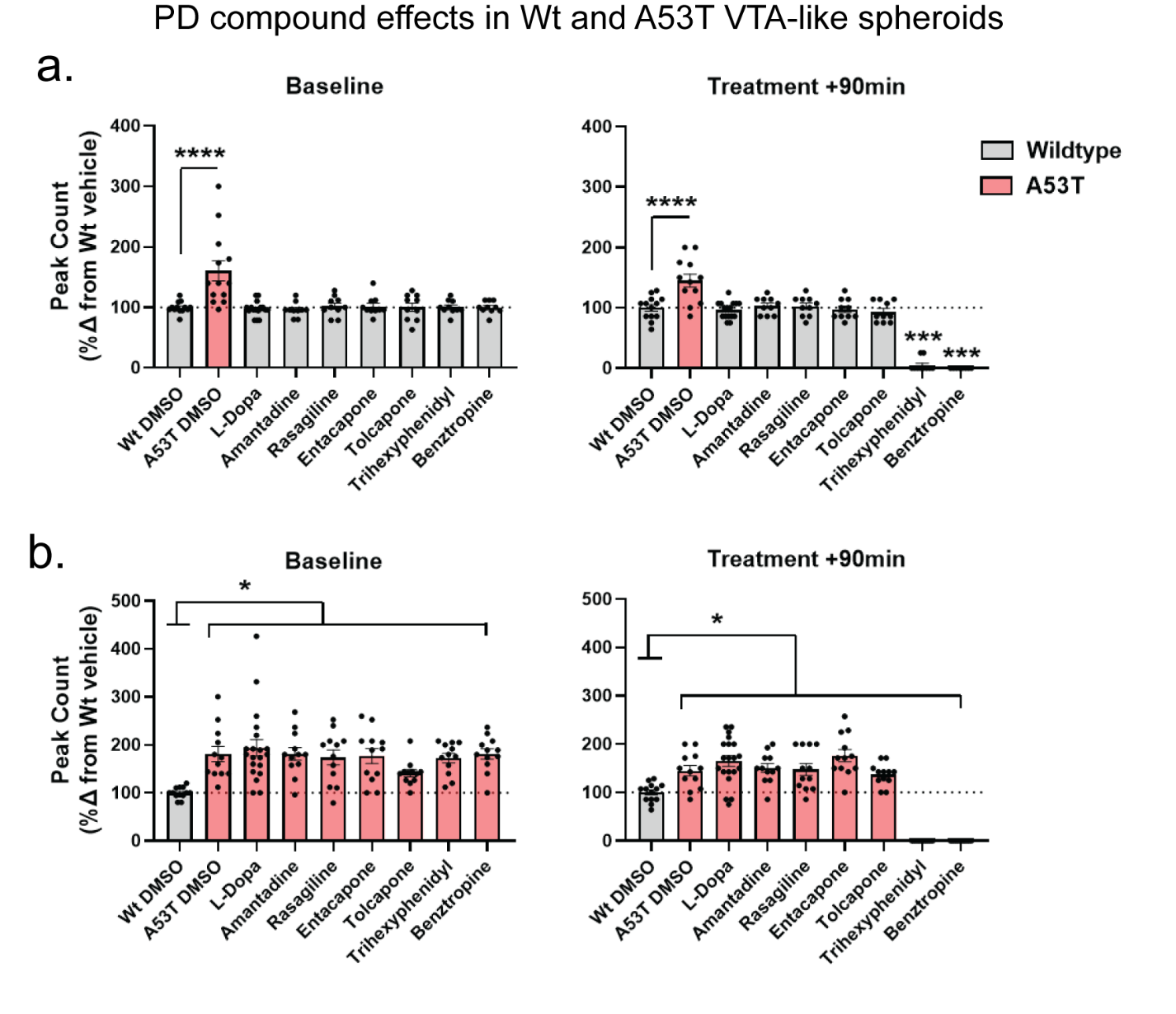
**Figure S5. Effects of compounds used to treat spheroids modeling Alzheimer’s Disease on Wt PFC-like spheroids. (a,b.)** Data collected from FLIPR recordings from spheroids incubating in Cal6 dye; Wt and APOE4 PFC-like spheroids at baseline and 90min after treatment with either DMSO or compounds used to treat AD. At baseline, significant chances in peak count **(a.)** and spacing **(b.)** were only observed in DMSO-treated APOE4 spheroids. At 90min, differences were observed in all groups except Rivastigmine (n=8-12 technical replicates, n=3 biological replicates collected over three independent experiments. Results are analyzed with One way ANOVA with significance set at p<0.05. Data from are represented as mean ± SEM; *p<0.05, **p<0.01, ***p<0.001, ****p<0.0001.

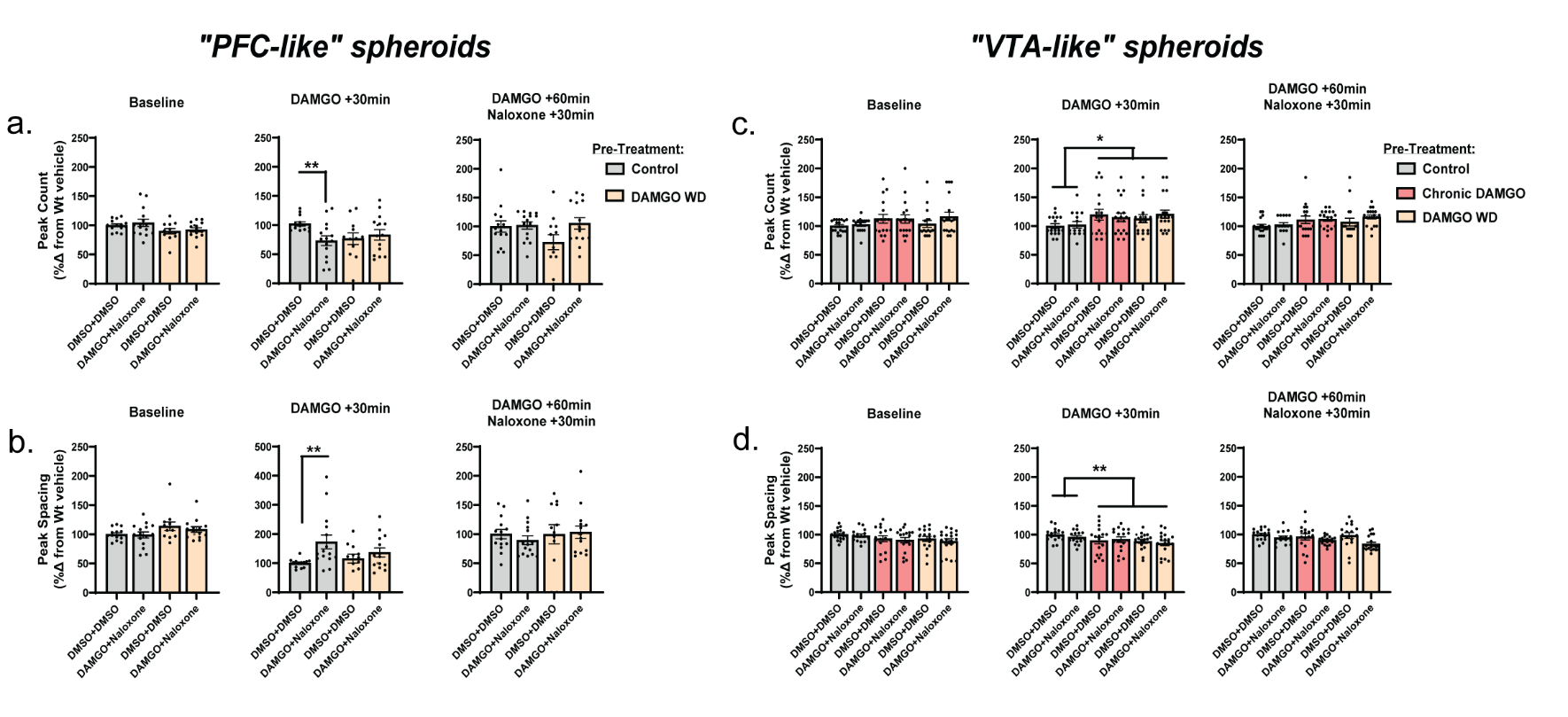
**Figure S6. Effects of compounds used to treat spheroids modeling Parkinson’s Disease on Wt and A53T VTA-like spheroids. (a,b.)** Data collected from FLIPR recordings from spheroids incubating in Cal6 dye; Wt and A53T VTA-like spheroids at baseline and 90min after treatment with either DMSO or compounds used to treat PD. **(a.)** Among Wt spheroids, significant chances in peak count were not observed at baseline. 90-min after treatment, significant changes in peak count were observed in treatment groups including Trihexyphenidyl and Benztropine **(b.)** At baseline and 90-min after treatment, A53T spheroids showed significantly increased peak count compared to Wt DMSO-treated controls across all treatment groups (n=8-12 technical replicates, n=3 biological replicates collected over three independent experiments. Results are analyzed with One way ANOVA with significance set at p<0.05. Data from are represented as mean ± SEM; *p<0.05, **p<0.01, ***p<0.001, ****p<0.0001.

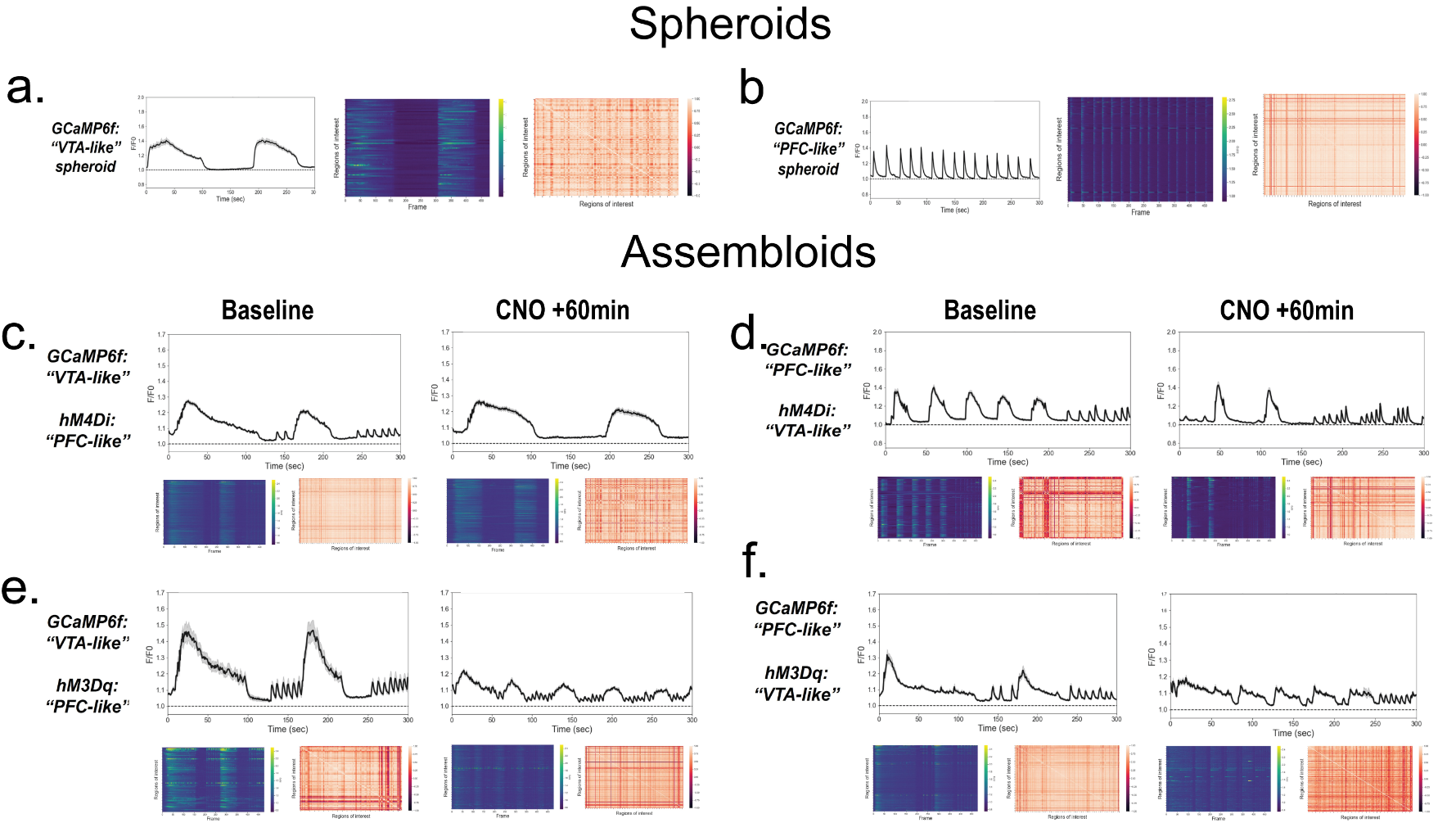
**Figure S7. Effects of naloxone on “PFC-like” and “VTA-like” spheroids modeling OUD. (a,b.)** Peak count **(a.)** and spacing **(b.)** in “PFC-like” spheroids modeling DAMGO WD compared to control spheroids receiving no pre-treatment. For both peak count and spacing, acute DAMGO treatment in control spheroids significantly changed peak count and spacing compared to DMSO-treated controls, and naloxone reversed these changes. No effects were observed in spheroids modeling DAMGO WD. **(c,d.)** Peak count **(c.)** and spacing **(d.)** in “VTA-like” spheroids. No effect of pre-treatment was observed during the baseline recording. 30-min after DAMGO or DMSO treatment, a main effect of pre-treatment was observed for both peak count and spacing. 30-min after naloxone or DMSO treatment, no effects were observed. (n=8-12 technical replicates, n=3 biological replicates collected over three independent experiments. Results are analyzed with Two way ANOVA with significance set at p<0.05. Data from are represented as mean ± SEM; *p<0.05, **p<0.01, ***p<0.001, ****p<0.0001.

**Figure S8. Phenotypic profiles in assembloids can be bidirectionally manipulated when one component of the assembloid through either chemogenetic stimulation or inhibition. (a,b)** Representative calcium activity from automated confocal calcium imaging recordings from a single “VTA-like” **(a.)** and “PFC-like” spheroid **(b.),** from left to right: average activity of all identified ROIs within spheroids, heatmap showing activity of all identified ROIs, correlation matrix indicating synchrony between all identified ROIs **(c-f.)** Representative calcium activity from automated confocal calcium imaging recordings from assembloids, which were made by fusing a “VTA-like” and “PFC-like” spheroid. Top: average activity of all identified ROIs within spheroids, lower left panel: heatmap showing activity of all identified ROIs, lower right panel: correlation matrix indicating synchrony between all identified ROIs. For each assembloid, activity is shown during the baseline recording as well as 60-min after clozapine-N-oxide (CNO) treatment to activate either stimulatory (hM3Dq) or inhibitory (hM4Di) DREADDs viruses **(c.)** Assembloid consisting of a “VTA-like” spheroid expressing GCaMP6f for calcium activity recordings, and a “PFC-like” spheroid expressing the inhibitory DREADDs virus, hM4Di **(d.)** Assembloid consisting of a “PFC-like” spheroid expressing GCaMP6f and a “VTA-like” spheroid expressing the inhibitory DREADDs virus, hM4Di **(e.)** Assembloid consisting of a “VTA-like” spheroid expressing GCaMP6f and a “PFC-like” spheroid expressing the stimulatory DREADDs virus, hM3Dq **(f.)** Assembloid consisting of a “PFC-like” spheroid expressing GCaMP6f and a “VTA-like” spheroid expressing the stimulatory DREADDs virus, hM3Dq

**Supplemental Table Legends**

**Table S1. Coefficients of variance calculated from 17 peak parameters extracted from ScreenWorks’ PeakPro 2.0 analysis.** Table showing percent coefficients of variance (%CV) values for 17 peak parameters obtained from peak analysis on calcium activity obtained on the FLIPR. %CV values were calculated by dividing the mean by the standard deviation then multiplying by 100 ((standard deviation/mean)*100) in Wt spheroids with no previous experimental manipulation. %CV values <30% indicated a peak parameter with low variability and those were used for future analysis and plotting.

**Table S2. Statistical analysis of functional responses to control compounds with 2-way repeated measures ANOVA.** Table representing p-values from multiple comparisons analysis with Sidak’s post hoc test comparing responses from each control compound to DMSO controls for each of the 10 peak parameters analyzed. Data was analyzed using linear mixed model ANOVA where treatment was a between-subjects factor and recording was a within-subjects factor to examine repeated measures. Significant treatment x recording interactions (p<0.05) were followed up with Sidak’s post hoc test, and p-values are displayed on the table, with those as bold being significantly different from DMSO.

**Supplemental Video Legends**

**Video S1. Expression of markers for neuronal subtype composition in a “VTA-like” spheroid.** Image stack showing expression of neuronal subtype markers within a “VTA-like” spheroid. From left to right, Hoechst nuclear stain, Tyrosine Hydroxylase (TH) for dopaminergic neurons, vGluT1 for glutamatergic neurons, Parvalbumin for GABAergic neurons. Z-stack was collected with a 0.5 µm step over 451 images using a 25X water objective.

**Video S2. Expression of markers for neuronal subtype composition in a “PFC-like” spheroid.** Image stack showing expression of neuronal subtype markers within a “PFC-like” spheroid. From left to right, Hoechst nuclear stain, Tyrosine Hydroxylase (TH) for dopaminergic neurons, vGluT1 for glutamatergic neurons, Parvalbumin for GABAergic neurons. Z-stack was collected with a 0.5 µm step over 439 images using a 25X water objective.

**Video S3. Calcium activity within a “VTA-like” spheroid. Automated confocal recording of calcium activity within a “VTA-like” spheroid.** Video was collected over 480 frames with a frame rate of 1.6 frames per second using a 20X objective.

**Video S4. Calcium activity within a “PFC-like” spheroid. Automated confocal recording of calcium activity within a “PFC-like” spheroid.** Video was collected over 480 frames with a frame rate of 1.6 frames per second using a 20X objective.
